# Estimation of neuronal firing rate using Bayesian Adaptive Kernel Smoother (BAKS)

**DOI:** 10.1101/204818

**Authors:** Nur Ahmadi, Timothy G. Constandinou, Christos-Savvas Bouganis

**Affiliations:** Centre for Bio-Inspired Technology, Institute of Biomedical Engineering, Imperial College London, London, SW7 2AZ, UK; Department of Electrical and Electronic Engineering, Imperial College London, London, SW7 2BT, UK

**Keywords:** Firing rate, kernel smoother, adaptive bandwidth, Bayesian framework, spike train model

## Abstract

Neurons use sequences of action potentials (spikes) to convey information across neuronal networks. In neurophysiology experiments, information about external stimuli or behavioral tasks has been frequently characterized in term of neuronal firing rate. The firing rate is conventionally estimated by averaging spiking responses across multiple similar experiments (or trials). However, there exist a number of applications in neuroscience research that require firing rate to be estimated on a single trial basis. Estimating firing rate from a single trial is a challenging problem and current state-of-the-art methods do not perform well. To address this issue, we develop a new method for estimating firing rate based on kernel smoothing technique that considers the bandwidth as a random variable with prior distribution that is adaptively updated under a Bayesian framework. By carefully selecting the prior distribution together with Gaussian kernel function, an analytical expression can be achieved for the kernel bandwidth. We refer to the proposed method as Bayesian Adaptive Kernel Smoother (BAKS). We evaluate the performance of BAKS using synthetic spike train data generated by biologically plausible models: inhomogeneous Gamma (IG) and inhomogeneous inverse Gaussian (IIG). We also apply BAKS to real spike train data from non-human primate (NHP) motor and visual cortex. We benchmark the proposed method against the established and previously reported methods. These include: optimized kernel smoother (OKS), variable kernel smoother (VKS), local polynomial fit (Locfit), and Bayesian adaptive regression splines (BARS). Results using both synthetic and real data demonstrate that the proposed method achieves better performance compared to competing methods. This suggests that the proposed method could be useful for understanding the encoding mechanism of neurons in cognitive-related tasks. The proposed method could also potentially improve the performance of brain-machine interface (BMI) decoder that relies on estimated firing rate as the input.

## 1. Introduction

In neural systems, signaling and interneuronal communication can be observed through the characteristic of action potentials (or ‘spikes’). A sequence of spikes, known as a spike train, may encode information based on different schemes. Currently, there are two main hypotheses of neural coding schemes: *temporal* coding and *rate* coding. The temporal coding represents the information by the precise timing or occurrence of spikes. On the other hand, the rate coding represents the information by the rate or frequency at which a neuron “fires” spikes, also known as “firing rate”, and has been the most commonly used scheme to characterize the neuronal or network responses to external stimuli or behavioral tasks [1,2]. The firing rate is typically estimated in offline analysis by averaging spiking responses across multiple repeated experiments known as trials. In practice, however, spiking responses may differ considerably even though the trial setting remains approximately the same. This is partly due to the inherent stochastic nature of neurons and the difference of cognitive states during the trials [3,4]. Averaging out many variably similar trials can obscure the temporal dynamics, which may contain useful information, on each single trial. Furthermore, many research of interests require the firing rate to be estimated on single trial basis. For example, quantifying trial-to-trial variability of neuronal responses [5,6], decoding task parameters in brain-machine interface (BMI) applications [3,5], and measuring neuronal responses in cognitive-related tasks such as decision making, motor planning, learning and memory [7–9]. Therefore, it is essential to be able to accurately estimate firing rate based on single trials.

Estimating firing rate as a continuous-time function from a single trial is a challenging task since the underlying process provides only a sparse representation of the spiking data. A widely used method known as peri-stimulus time histogram (PSTH) results in a coarse estimate [10,11]. To produce smooth estimate of firing rate, several methods have been proposed such as optimized kernel smoother (OKS) [12], variable kernel smoother (VKS) [12], local polynomial fit (Locfit) [13], and Bayesian adaptive regression splines (BARS) [14]. The OKS and VKS methods employ kernel density estimation technique in which the accuracy of estimation is heavily impacted by the choice of the kernel bandwidth. Both OKS and VKS methods automatically compute the bandwidth based on mean integrated squared error (MISE) minimization principle. However, in computing the bandwidth, these methods make assumption that spikes are generated by a Poisson process. Even though superimposed spike trains across many trials approximate a Poisson process, in the case of single trial, a spike train has been shown to depart from this assumption [15,16]. Single trial spike train exhibits history-dependent properties such as refractory period and bursting, which cannot be modeled by Poisson process [15,16]. The deviation from Poisson assumption can lead both OKS and VKS methods to exhibit poor performance under single trial cases. Locfit method employs generalized nonparametric regression technique where the firing rate is approximated by polynomials. The estimation accuracy of Locfit depends mostly on a smoothing parameter (bandwidth). How this bandwidth is selected along with the Poisson assumption of the spike train are the main caveat of this method. Like Locfit, BARS also employs generalized regression technique, except that it estimates the firing rate using splines (several polynomials connected at some points or knots). The challenge of using a spline-based method is determining the number and location of the knots since these will significantly impact the estimation. To determine the optimal knot configurations (number and location), BARS utilizes reversible-jump Markov chain Monte Carlo (MCMC) engine with Bayesian information criterion (BIC). The flexibility and powerfulness of BARS comes at a price of relatively high computational complexity [14,17]. In addition, similarly to above-mentioned methods, BARS also assumes that spikes are generated from a Poisson process. It is intended to be used for firing rate estimation after pooling spikes across multiple trials [17].

Within this context, it is highly desirable to have a method for estimating neuronal firing rate from a single trial spike train, which features an adaptive capability and low computational complexity. An adaptive capability is crucial as spiking activity may change rapidly within single trial. The proposed method should be data-driven and is able to accurately estimate the underlying spike dynamics. Low computational complexity is needed to perform the computation within a short time, and is necessary for real-time BMI applications. In addition, it is important to have a spike train model that mimics the spiking behavior encountered in real neural recording. Using certain assumption on the underlying rate function (i.e. the “ground truth” is known), single spike trains can be stochastically generated which can then be used to evaluate the performance of the firing rate estimation method.

In this paper, we propose a new method for estimation of firing rate that addresses the issues listed above. This method employs kernel smoothing technique due to its fast and simplicity. The key parameter, bandwidth, is considered as a random variable with a prior distribution and is adaptively determined under a Bayesian framework. With the appropriate selection of kernel and prior distribution functions, analytical expression of posterior distribution can be attained which reduce the computational complexity. We refer to this method as Bayesian Adaptive Kernel Smoother (BAKS). We evaluate the proposed method with synthetic data generated from two biologically plausible models: inhomogeneous Gamma (IG) and inhomogeneous inverse Gaussian (IIG). The proposed method is then tested with real neural data recorded from motor and visual cortex of non-human primate (NHP).

## 2. Methods

In this section, we first introduce a kernel smoothing technique for estimating firing rate. We then describe our proposed method, BAKS, a new variant of kernel-based firing rate estimation method that incorporates adaptive bandwidth. Lastly, we explain two models that we used to generate synthetic spike train data for evaluating the performance of the proposed method. The BAKS code and all the datasets that we synthesized (in Matlab) have been made publicly available through https://github.com/nurahmadi/BAKS.

### 2.1. Kernel-based firing rate estimation

Let *t*_1_, *t*_2_, … , *t_n_* be a sequence of spike times (i.e. a spike train) which can be expressed mathematically as

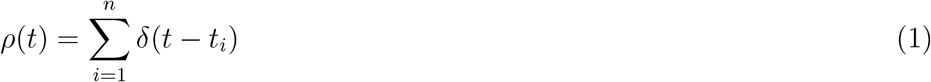

where *δ*(*t*) is Diract function and *n* is the total number of spikes. The underlying rate function also known as firing rate, λ(*t*), can be estimated by using kernel smoothing, a method which convolves the spike train with a kernel function *K*(*t*) as follows,

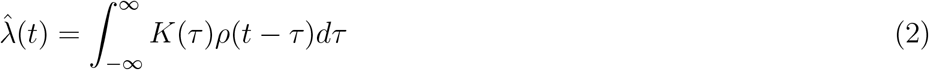

Eq (2) can also be represented as the sum over kernel function centered at spike times *t_i_*,

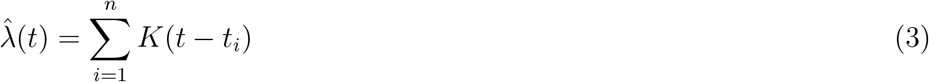

It has been suggested that the choice of kernel function is not critical to the estimation result [5,18]. Any non-negative kernel function that is normalized to a unit area, satisfies a zero first moment, and has a finite variance can be used for firing rate estimation [5,12]. Among the several choices, the Gaussian is the most frequently used kernel function and is defined by

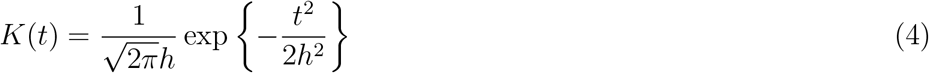

where *h* > 0 is the bandwidth (kernel width), the main parameter that controls the smoothness of the estimate. In Eq (4), the bandwidth is fixed over the whole observation interval. A significant amount of literature has been reported in the field of statistics on selecting the proper value for this fixed bandwidth [19–23]. Even though a near-optimal fixed bandwidth selection (e.g. [12,24,25]) may yield a better estimate compared to an arbitrary choice, it may still suffer from simultaneously under-and over-smoothing depending on the underlying spike dynamics. A rapid change in spiking activity is sometimes encountered in neural responses and is of interest to neuroscientists. Thus, it is highly desirable to find optimal adaptive bandwidth selection method that can adaptively grasp the slow and rapid changes of firing rate.

### 2.2. Bayesian Adaptive Kernel Smoother (BAKS)

BAKS incorporates adaptive bandwidth at the estimation points, meaning the bandwidth at which firing rate is being estimated can adapt to the dynamics of the underlying process. A Gaussian kernel with adaptive bandwidth is expressed as follows,

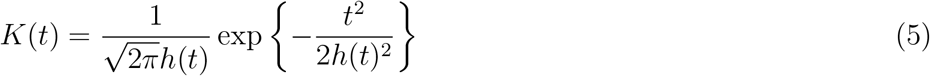

where *h*(*t*) is the adaptive bandwidth. The bandwidth should be small (large) at the region of high (low) spike density.

To find the optimal adaptive bandwidth, BAKS considers the bandwidth as a random variable with prior distribution and adaptively updates the posterior bandwidth given spiking data using Bayes’ theorem. We select a prior distribution of bandwidth that incorporates prior belief about spiking data (i.e. informative prior) and leads to an analytical expression of posterior distribution. The informative prior is especially useful when the number of spiking data is relatively small as this prior will give more weight than the likelihood function; while the analytical expression is of paramount importance to simplifying the process and avoiding having computational burden of posterior sampling techniques (e.g. Markov chain Monte Carlo). In Eq (4), the Gaussian kernel uses parameter bandwidth (i.e. standard deviation in statistical literature) that describes how spread the observed data are around the mean. We can also represent the parameter in term of precision (inverse of square bandwidth) that describes how concentrated the observed data are around the mean. In computing firing rate estimation using Eq (3), the means of Gaussian kernel are set to the spike times. These spike times can be represented as sum of the interspike intervals (ISIs) which according to a number of research can be conveniently modeled by Gamma distribution [15,26–29]. Since sum of independent Gamma random variables follows Gamma distribution [30], the spike times can also be represented as Gamma distribution. Hence, we propose Gamma prior distribution on the precision parameter (σ), where σ = 1/*h*(*t*)^2^. As Gamma distribution is a conjugate prior for Gaussian distribution with precision parameter [31], the choice of Gamma prior distribution results in closed-form expression of the posterior distribution. This Gamma prior distribution is given by

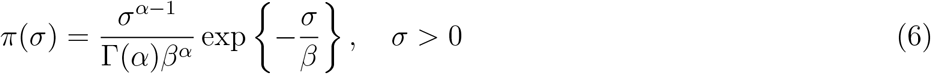

where *α* > 0 is the shape parameter, *β* > 0 is the scale parameter, and Γ(α) is Gamma function. By the change-of-variable formula and transformation technique, we can express the prior distribution π(σ) as a function of *h*(*t*):

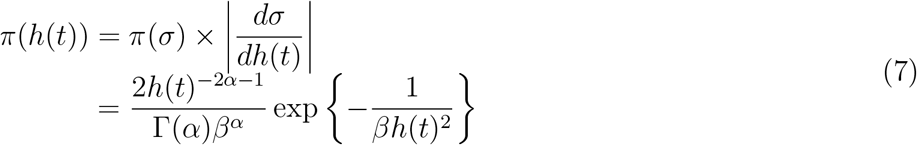

The likelihood function of bandwidth given the spike train is equal to probability density of spike train conditional to bandwidth parameter, which can be approximated by

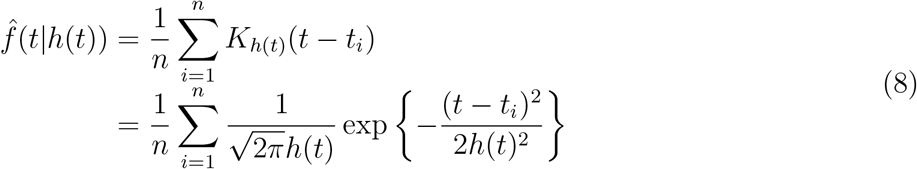

where *K*_*h*(*t*)_(*t* - *t_i_*) is Gaussian kernel as defined in Eq (5) with adaptive bandwidth and centered at spike event time *t_i_*.

Using Bayes’ theorem, the posterior distribution of kernel bandwidth, π(*h*(*t*)|*t*), can be computed by

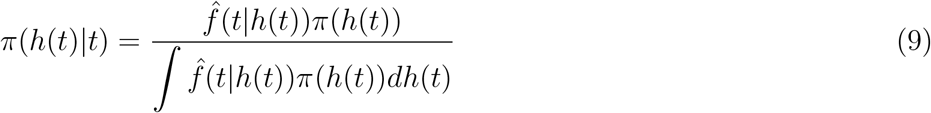

The denominator also called the normalizing factor can be problematic if there is no analytical solution of its integral expression since a burdensome numerical approximation method will be required instead. The choice of prior distribution in Eq (7) coupled with Gaussian kernel density in Eq (8) leads to an analytical expression for the denominator in Eq (9) as follows,

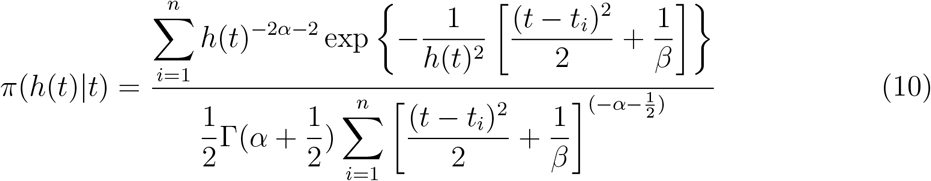

The adaptive bandwidth can then be estimated under squared error loss function by using the posterior mean formulated as

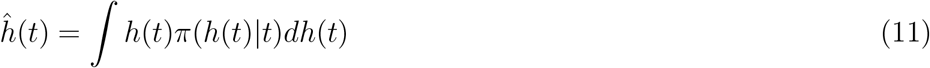

The closed-form expression of Eq (11) is given by

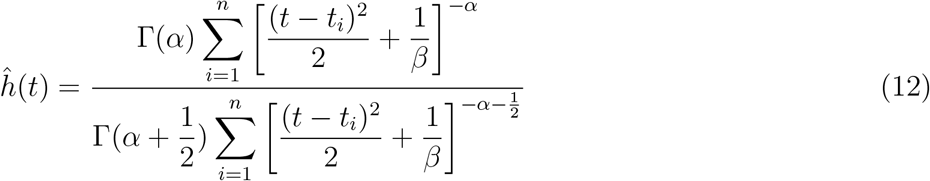

The derived bandwidth is then used in firing rate estimation formula given by Eq (3). The BAKS method is graphically illustrated in Fig 1. The derivation of the closed-form expressions of bandwidth posterior distribution and the adaptive bandwidth estimate are given in Appendix A and Appendix B, respectively.

#### 2.2.1. Selection of prior parameters

The accuracy of firing rate estimation depends on the bandwidth parameter, which is influenced by the selection of parameter *α* and *β* of the prior distribution. According to bandwidth prior distribution in Eq (7), we can calculate its mean and variance as follows,

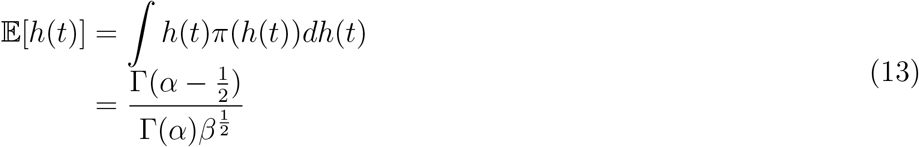

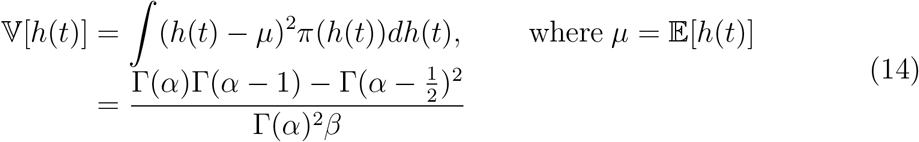

where 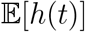 and 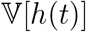 are the mean and variance of π(*h*(*t*)), respectively.

**Figure 1.**
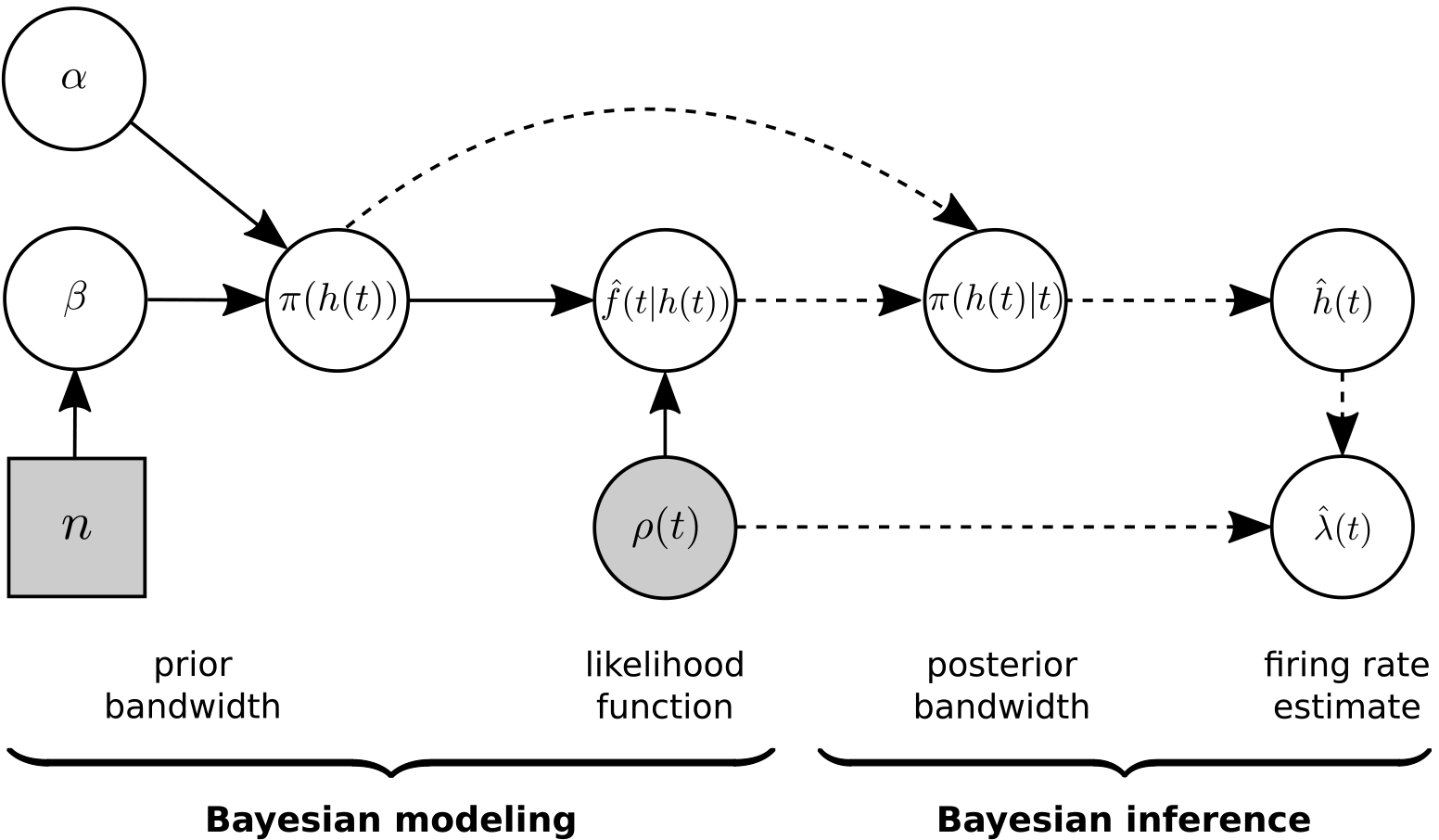
Graphical representation of the BAKS model. The various blocks describe the different components as follows: Circular blocks denote continuous variables, and rectangle blocks denote discrete variables. Shaded blocks represent observed variables, whereas white blocks represent hidden variables. Solid and dashed arrow indicate Bayesian modeling and inference phase, respectively.

It can be observed from both Eqs (13) and (14) that the mean and variance are inversely proportional to the value of *β*. As the number of spike events (*n*) increases, the bandwidth decreases. Hence, to obtain consistency of the estimation, choice of *β* is proposed to be a function proportional to the number of spike events (*n*). Here, we set *β* = *n*^4/5^ in accordance with MISE convergence rate of Gaussian kernel [32].

Since *h*(*t*) > 0, the numerator of Eqs (13) and (14) must be greater than zero, which in turn requires *α* > 1. The optimal value of *α* can be determined by minimizing MISE function given fixed value of *β* as follows,

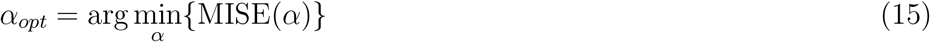

MISE is a common metric used in evaluating goodness-of-fit of a density estimation and has been used in previous studies of firing rate estimation [5,12,33]. MISE is computed between the estimated firing rate 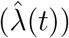 and the true underlying rate function (*λ*(*t*)) with the following formula,

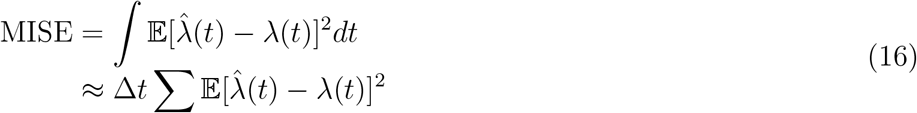

where 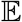 denotes the expectation with respect to stochastic process of spike generation model and the integration/summation is performed over observation interval.

### 2.3. Spike train generation model

It is essential to have a model for generating spike trains in order to train the firing rate estimation method (i.e. find the optimal parameter) and for evaluating performance. A spike train can be modeled by using a point process; a stochastic process that describes localized events or points in real (e.g. time, space) domain. The most commonly used class of point processes to model a spike train is inhomogeneous Poisson (IP) process. [3,5,12,15,24]. In this class, each spike time is assumed to be independent or memoryless, meaning that a spike occurring at a particular time does not depend on the past spiking activities. It has been shown that IP process can be used to approximate the behavior of neural spikes when the spikes are superimposed across trials [12,16]. However, a Poisson process cannot model the history-dependent properties (e.g. refractory period and bursting) of neural spike trains in the case of single trial. Hence, another class of point process is required to model single trial spike train.

*The renewal process* is an alternative class of point process that can describe the history-dependent property of spike times by assuming that a spike fired at any point in time depending only on the last spike, not the spikes before it. In the renewal process, spike times are no longer independent, but it is their inter-spike intervals (ISIs) that remain independent. Spike trains can be conveniently represented by their ISIs drawn from a certain distribution. Two classes of distribution which have often been used to model non-Poisson spike train are Gamma and inverse Gaussian distributions [6,15,26–29,34]. The former is then called inhomogeneous Gamma (IG) model while the latter is called inhomogeneous inverse Gaussian (IIG) model.

#### 2.3.1. Inhomogeneous gamma (IG) model

The Inhomogeneous Gamma (IG) model generalizes a Poisson model by allowing more flexible ISI distribution that is controlled by shape parameter (γ) [15,26–29]. The gamma probability density of the ISI is defined as

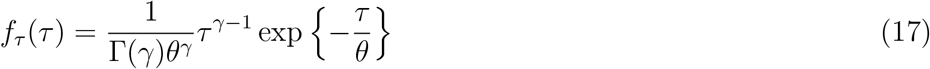

where τ > 0 denotes the ISI, γ > 0 represents the shape parameter, *θ* is the scale parameter, and Γ(γ) is the Gamma function. As shown in Eq (17), when γ =1, the ISI becomes exponentially distributed known as Poisson model. If γ < 1, the probability density value declines faster than exponential, which means the ISI tends to become smaller. This can be used to describe neuron’s rapid firing (bursting) phenomena. If γ > 1, the probability density value will increase from zero up to a certain peak value and then decrease again to zero. This indicates the refractory period property where a neuron is less likely to fire again immediately after it fires a spike.

To generate spike train from such a model, we employ time-rescaling theorem which transforms the original spike times into new rescaled spike times where the ISIs are independently and identically drawn from fixed distribution (e.g. Gamma) [28]. In the IG model, this transformation is defined as integration of underlying rate function, λ(*t*), on the interval (0, *t*] and multiplied by the shape parameter (γ). This transformation is formulated as

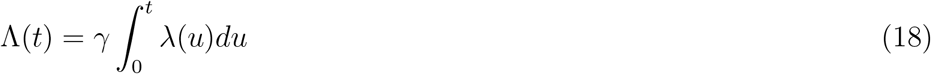

Since λ(*t*) is a non-negative function, ^(*t*) is then a monotonically increasing and one-to-one function. Thus, we can obtain the original spike times from *i.i.d.* ISI samples (τ) by performing the inverse of time-rescaling transform as follows,

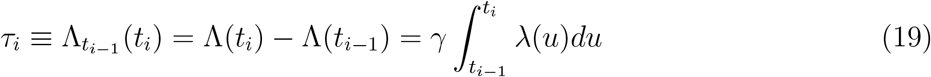

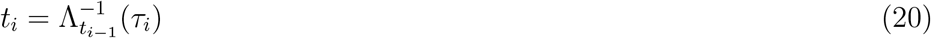

where 0 < *t*_1_ < *t*_2_, … , *t_n_* ≤ *T* represent the spike times within interval (0, *T*] and 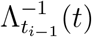 is the inverse of time-rescaling transform.

#### 2.3.2. Inhomogeneous inverse Gaussian (IIG) model

It has been suggested that inhomogeneous inverse Gaussian (IIG) model is more biologically plausible to describe the characteristic of neural spike train than IG model [15,35]. The IIG model uses inverse Gaussian distribution for the ISI and has been applied in multiple studies [36,37]. The inverse Gaussian ISI distribution used in this model is given by

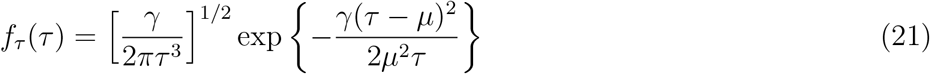

where τ > 0 denotes the ISI, γ is the shape parameter, and *μ* is the location parameter. In the IIG model, the value of ISI density is zero at the origin, then increases to certain peak value, and decreases again to zero. How quick the density value rises or falls is controlled by the shape parameter (γ), which demonstrates the flexibility of IIG model in describing different neural spike characteristics such as rapid bursting and refractory period. The smaller (larger) value of γ, neuron tends to fire more bursty (regularly).

To generate a spike train using the IIG model, we employ a similar procedure as that of IG model, with the exception that the time-rescaling transform here is expressed as

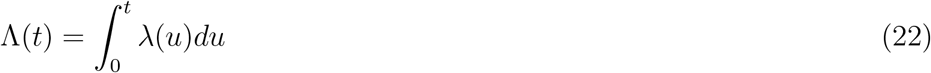

where the interval between subsequent spike times (ISIs) in the rescaled domain are drawn from *i.i.d.* inverse Gaussian distribution. The spike times in the original domain can be computed by using the inverse of time-rescaling transform as shown in Eq (20).

## 3. Results

### 3.1. Synthetic datasets

In our synthetic data, the spike trains were generated from IG and IIG models with different underlying rate functions that represent non-stationary processes usually encountered in empirical datasets. These rate functions include (1) a continuous process with changing (heterogeneous) frequency, (2) a continuous process with oscillatory (homogeneous) frequency, and (3) a discontinuous process with sudden rate changes. In this study, the first, second, and third rate functions are referred to as ‘chirp’, ‘sine’, and ‘sawtooth’ rate functions, and are mathematically expressed as,

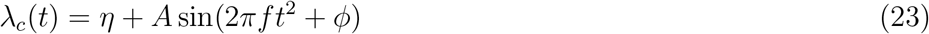

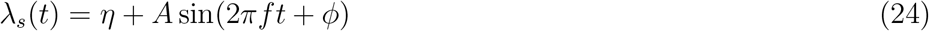

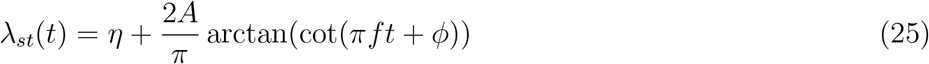

where *η* indicates the base or average number of spikes per second and *A* denotes the intensity (or amplitude) which controls the dynamic range of the rate function. Parameter *f* is frequency whereas *ϕ* is phase. The chirp, sine and sawtooth rate functions are represented by *λ_c_*(*t*), λ_*s*_(*t*) and λ_*st*_(*t*), respectively.

The chirp rate function used in this study is similar to that of Rao and Teh [38], whereas the sine and sawtooth processes are similar to that of Shimazaki and Shinomoto [12] but with different intensity, frequency, and spike train model. Examples of chirp, sine, and sawtooth rate functions with certain parameter settings are illustrated in Figs 2a, 2b, 2c, respectively.

During parameter tuning of BAKS (training phase), we set *η* to 50 spikes/s and *A* to 25 spikes/s (corresponds to dynamic intensity range between 25 and 75 spikes/s) as in [27]. These intensity values are referred to as medium intensity. This setting is reasonable as in practice cortical neurons do not often fire more than 100 spikes/s [39]. The frequency of chirp (*f_c_*), sine (*f_s_*) and sawtooth (*f_st_*) rate functions were set to 0.5, 1 and 1, respectively. These frequency values are referred to as medium frequency. We generated randomly 100 single spike trains (duration of 2 s) from two models (IG and IIG) and three rate functions (chirp, sine, and sawtooth). In IG model, the shape parameter (γ) that determines the regularity of firing rate was set to 4, while the scale parameter (*θ*) was set to 1. This setting is comparable to real neural spiking data and also used in [26,34]. Analogously, the shape (γ) and location (*μ*) parameters in IIG model were respectively set to 4 and 1. The optimal parameter value of prior distribution in BAKS method (*α*_*opt*_) was selected by minimizing the average MISE from two models and three rate functions mentioned above. The same procedure was performed to find the optimal smoothing parameter for Locfit.

For evaluating the performance of BAKS (testing phase), we generated 5 datasets each containing 100 repetitions from both IG and IIG models with the same and different setting from that of training phase. Dataset testing 1 was generated using the same setting as in the training phase. Dataset testing 1 was used to evaluate the BAKS when the underlying rate functions are the same but their stochastic spiking data realizations are different. Dataset testing 2 was used to evaluate the proposed method when the frequency and intensity of the underlying rate functions are different. The frequency of chirp, sine and sawtooth rate functions were respectively set to low (0.25, 0.5 and 0.5) and high (0.75, 1.5 and 1.5) while keeping the intensity same as that of training phase. Then, the dynamic intensity range of these three rate functions were set to low (2 - 20) and high (25 - 175) while keeping the frequency same as that of training phase. In dataset testing 3, the model setting was equal to during the training except that the parameter shape (γ) was varied between 1 and 10 with an increment of 1. Dataset testing 4 was used to assess the performance of BAKS under multiple trials (*tr* = {5,10, 20, 30}). These multi trial spike trains were obtained by superimposing a number of trial (5, 10, 20 or 30) into a single spike trains with the same setting as of training. In dataset testing 5, spike trains were generated using Gaussian-damped sinusoidal rate function with variable intensity and frequency. The idea of using this rate is taken from Mazurek’s work [39]. Mathematically, the Gaussian damped sinusoidal rate function is given by

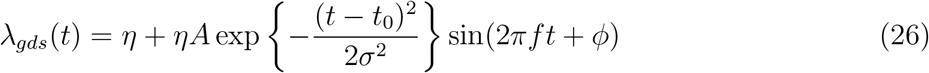

**Figure 2.**
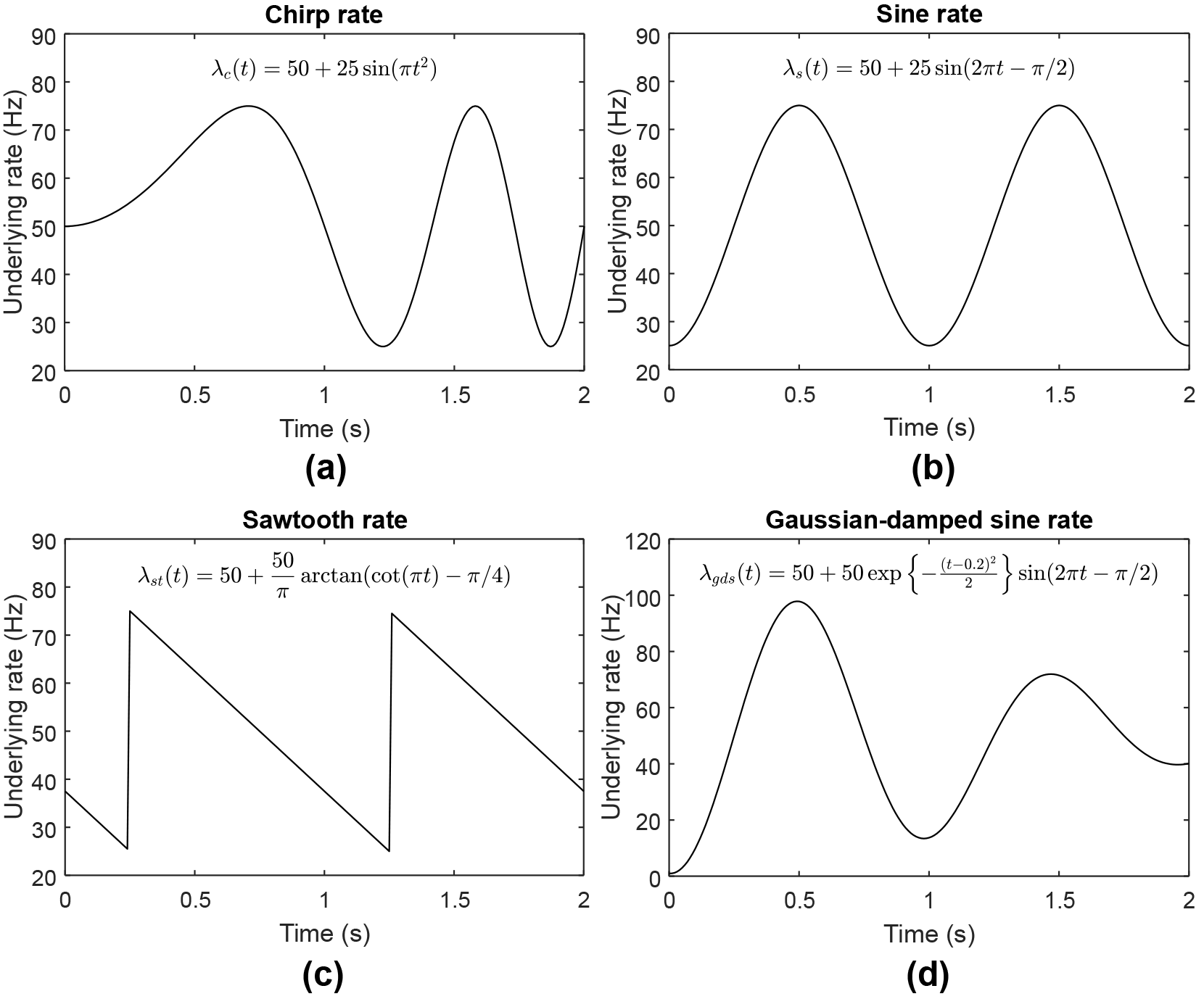
Illustration of the underlying rate functions used in this study. (a) Chirp rate function with η = 50, *A* = 25, *f_c_* = 0.5, and *ϕ* = 0. (b) Sine rate function with η = 50, *A* = 25, *f_s_* = 1, and ϕ = -π/2. (c) Sawtooth rate function with η = 50, *A* = 25, *f_st_* = 1, and ϕ = -π/4. (d) Gaussian-damped sinusoidal rate function with η = 50, *A* =1, *t*_0_ = 0.2, *σ* =1, *f_gds_* = 0.5, and ϕ = -π/2.

During the generation of dataset testing 5, the frequency (*f_gds_*) was set to low (0.5), medium (1.5) and high (2) while keeping the dynamic intensity range to medium (0 to 100) which corresponds to η = 50 and *A* =1; the dynamic intensity range was varied between low (0 to 20), medium (0 to 100) and high (0 to 200) while setting *f_gds_* = 1. An example of Gaussian-damped sinusoidal rate function is illustrated in Figs 2d. The summary of model setting for generating the synthetic datasets for training and testing is shown in Table 1.

**Table 1.** Model setting for synthetic dataset generation.

| Phase | Rate function | Intensity | Frequency | Parameter shape ($\gamma$) | Number of trial ($tr$) |
| --- | --- | --- | --- | --- | --- |
| Training | {Chirp, sine, sawtooth} | Medium | Medium | 4 | 1 |
| Testing 1 | {Chirp, sine, sawtooth} | Medium | Medium | 4 | 1 |
| Testing 2 | {Chirp, sine, sawtooth} | {Low, high} | {Low, high} | 4 | 1 |
| Testing 3 | {Chirp, sine, sawtooth} | Medium | Medium | {1, 2, …, 10} | 1 |
| Testing 4 | {Chirp, sine, sawtooth} | Medium | Medium | 4 | {5, 10, 20, 30} |
| Testing 5 | {Gaussian-damped sine} | {Low, medium, high} | {Low, medium, high} | 4 | 1 |

Training: medium intensity ( $\eta = 50, A = 25$ ), medium frequency ( $f_c = 0.5, f_s = 1, f_{st} = 1$ )
Testing 1: medium intensity ( $\eta = 50, A = 25$ ), medium frequency ( $f_c = 0.5, f_s = 1, f_{st} = 1$ )
Testing 2: low intensity ( $\eta = 11, A = 9$ ), high intensity ( $\eta = 100, A = 75$ ), low frequency ( $f_c = 0.25, f_s = 0.5, f_{st} = 0.5$ ), high frequency ( $f_c = 0.75, f_s = 1.5, f_{st} = 1.5$ )
Testing 3: medium intensity ( $\eta = 50, A = 25$ ), medium frequency ( $f_c = 0.5, f_s = 1, f_{st} = 1$ )
Testing 4: medium intensity ( $\eta = 50, A = 25$ ), medium frequency ( $f_c = 0.5, f_s = 1, f_{st} = 1$ )
Testing 5: low intensity ( $\eta = 10, A = 1$ ), medium intensity ( $\eta = 50, A = 1$ ), high intensity ( $\eta = 100, A = 1$ ), low frequency ( $f_{gds} = 0.5$ ), medium frequency ( $f_{gds} = 1.5$ ), high frequency ( $f_{gds} = 2$ )

### 3.2. Selection of optimal prior parameter (α)

We investigated the choice of parameter value that yielded the most accurate firing rate estimates measured by MISE metric. The optimal parameter (*α_opt_*) obtained from the training phase was then used as parameter values for our proposed method during testing phase. Spike trains used during training phase were stochastically generated from two models (IG and IIG) and three different underlying rate functions (chirp, sine, sawtooth) as described in previous section. We performed firing rate estimation over training dataset using parameter α varying from 1 to 10 with an increment of 0.5. This estimation was carried out while keeping parameter *β* fixed to *n*^4/5^, where *n* represents total number of spikes within observed duration.

The performance of estimation was quantified in term of MISE, mean of integrated squared error between the underlying and estimated firing rates over the observed duration (2 s). From 100 repetitions and 19 variation of parameter values (*α* = {1,1.5, 2, …, 10}), we computed the MISE values along with their 95% confidence interval from these 6 scenarios (2 models and 3 rate functions). The MISE values and their confidence interval for IG and IIG models are shown in Figs 3a and 3b. In IG model, *α* value that resulted in smallest MISE for chirp, sine and sawtooth rate function are 4, 3 and 6, respectively. In IIG model, α value associated with the smallest MISE for chirp, sine and sawtooth rate function are 4.5, 3 and 5.5, respectively. To find the optimal parameter, we computed average MISE values across 6 scenarios. The value of α that corresponds to the smallest average MISE was then determined as the optimal parameter. According to the results as shown in Fig 3c, we selected 4 as the optimal parameter (*α_opt_*) and used this during the testing phase.

**Figure 3.**
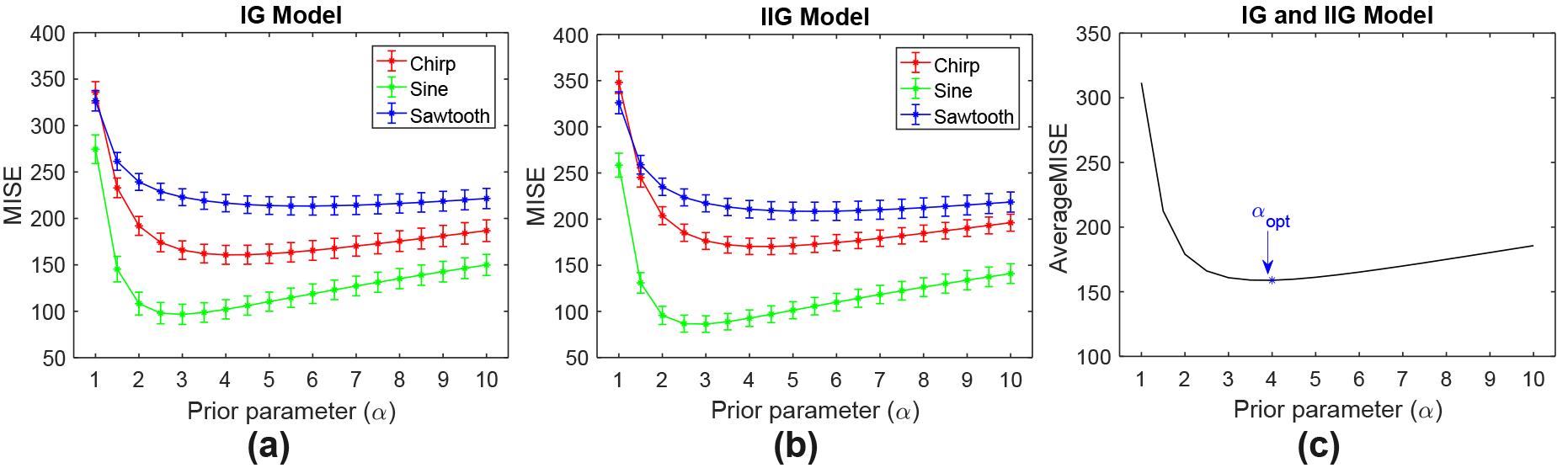
MISE comparison for various prior parameter (α) values. (a) MISE values (star mark) and their 95% confidence interval (vertical bar) for IG model. (b) MISE values (star mark) and their 95% confidence interval (vertical bar) for IIG model. (c) Average MISE values across 2 models and 3 rate functions. Optimal parameter *α_opt_* = 4 was selected as it yielded the smallest average MISE.

### 3.3. Comparison with the established methods

We evaluated and compared the performance of the proposed method (BAKS) with the established methods which include optimized kernel smoother (OKS) [12], variable kernel smoother (VKS) [12], local polynomial fit (Locfit) [13] and Bayesian adaptive regression splines (BARS) [14]. The performances were quantified using MISE as expressed in Eq (16). We did not include the the histogram (PSTH) method in the comparison as this cannot produce a smooth estimate under single-trial case. Shimazaki and Shinomoto demonstrated that even if the number of trials is increased, the performance of PSTH is far outperformed by OKS, VKS, Locfit, and BARS [12].

The two kernel smoothing methods used in the presented work, OKS and VKS, were developed by Shimazaki and Shinomoto [12]. In the OKS method, the bandwidth is fixed for the whole duration and automatically selected based on global MISE minimization principle. Unlike OKS, the VKS method employs variable bandwidth and this bandwidth is automatically determined by minimizing local MISE function. Thus, both
OKS and VKS methods do not require manual user intervention in selecting optimal bandwidth parameter. The Matlab codes for OKS and VKS methods can be obtained from the author’s website (http://www.neuralengine.org/res/kernel.html).

The Locfit is part of Chronux analysis software that at the time of this study can be downloaded from http://chronux.org/. This method estimates the firing rate by maximizing local log-likelihood where the log-density function is approximated by local polynomial. The Locfit has some parameters such as degree of polynomial, weight function, and bandwidth. However, bandwidth is considered as the most crucial parameter that affects the accuracy of estimation [13]. Therefore, in Locfit setting, we concerned more on the bandwidth selection than other parameters. We used nearest neighbor bandwidth so that the local neighborhood always contains sufficient data (spikes). This can reduce data sparsity problem that may arise in real neural data. The nearest neighbor bandwidth parameter was determined by training procedure, while the parameter of degree of polynomial and weight function were set to the default values which are two and tricube (‘tcub’), respectively. The training procedure in Locfit was similar to that of α parameter in our proposed method. We performed MISE comparison from three underlying rate processes (chirp, sine, and sawtooth) for nearest neighbor bandwidth between 0.2 to 0.8 with increment 0.1. We chose 0.4 as the optimal parameter value since the average MISE associated with this value was the smallest among others. The value 0.4 means that the Locfit uses 40% of the total data in each estimation point.

The BARS method estimates the firing rate by using cubic spline basis function with free parameters on the number and location of knots [14]. The optimal knot configurations is determined by a fully Bayesian approach with reversible-jump Markov chain Monte Carlo (MCMC) engine and locality heuristic. The BARS takes spike count within bin interval (histogram) centered on estimation time of interest. In our study, we used Matlab implementation of the BARS which is available at http://lib.stat.cmu.edu/~kass/bars/bars.html. We used Poisson prior distribution of the knots and set the sample iterations to 5000 and burn-in samples to 500. We used spike count within 10 ms bin interval as the input and mean of fitted function as the output estimate.

We performed 100 repetitions of single-trial firing rate estimation using dataset testing 1. We plotted the MISE of all the methods and their confidence interval for three underlying rate functions and two models (Fig 4). In all 6 scenarios, the MISE and confidence interval of the BAKS method were smallest among other competing methods. Clearly, as shown in Fig 4a for IG model and Fig 4b for IIG model, the BAKS method (blue bar) yielded significantly lower MISE compared to the other competing methods. This demonstrates the effectiveness of the proposed method in estimating the underlying rate from single trial spike trains.

Examples of the underlying rate functions and the estimated firing rates from all methods across 6 scenarios are shown in Fig 5. In the cases of chirp and sawtooth underlying rate processes, where the frequency of these function (thus the spike density) change over the observed duration, the BAKS and VKS methods which feature adaptive or variable bandwidth, appear visually better compared to the rest methods (see Figs 5a and 5c, and Figs 5d and 5f). This indicates the reliability of the adaptive or variable bandwidth employed by the two methods in adapting to different density over the duration. On the other hand, OKS and Locfit which use fixed bandwidth cannot produce good fit for the chirp and sawtooth cases. The BARS method, despiting having an adaptive capability, yields poor estimates. This may be due to the insufficient number of spikes (data) available in single trial required by the BARS to produce good estimates. However, in the case of sine underlying rate process (homogeneous frequency), the goodness-of-fit across all methods does not seem significantly different (Figs 5b and 5d).

**Figure 4.**
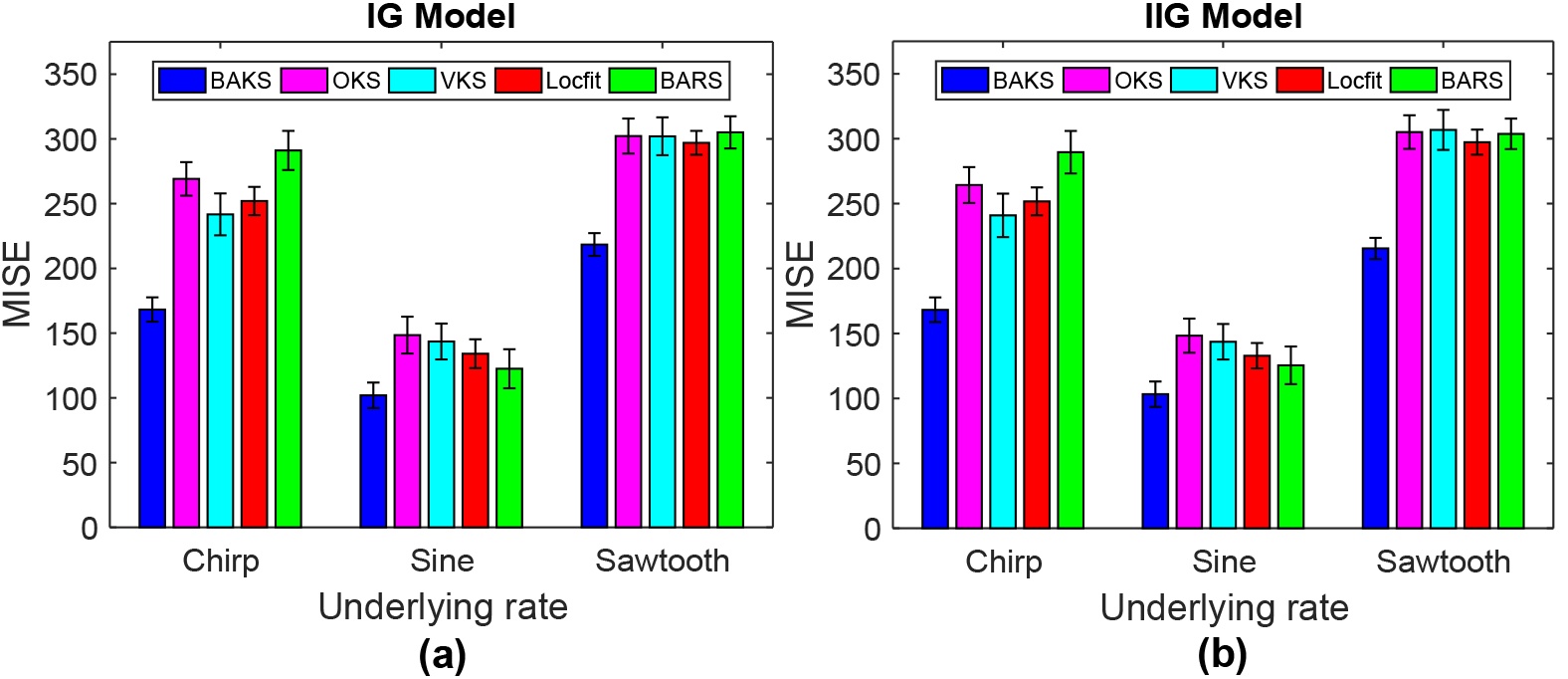
MISE comparison of BAKS with other methods. (a) MISE comparison for three rate processes (chirp, sine, and sawtooth) from IG model. (b) MISE comparison for three rate processes (chirp, sine, and sawtooth) from IIG model. Vertical line crossing the peak of bar plot represents the 95% of MISE confidence interval.

**Figure 5.**
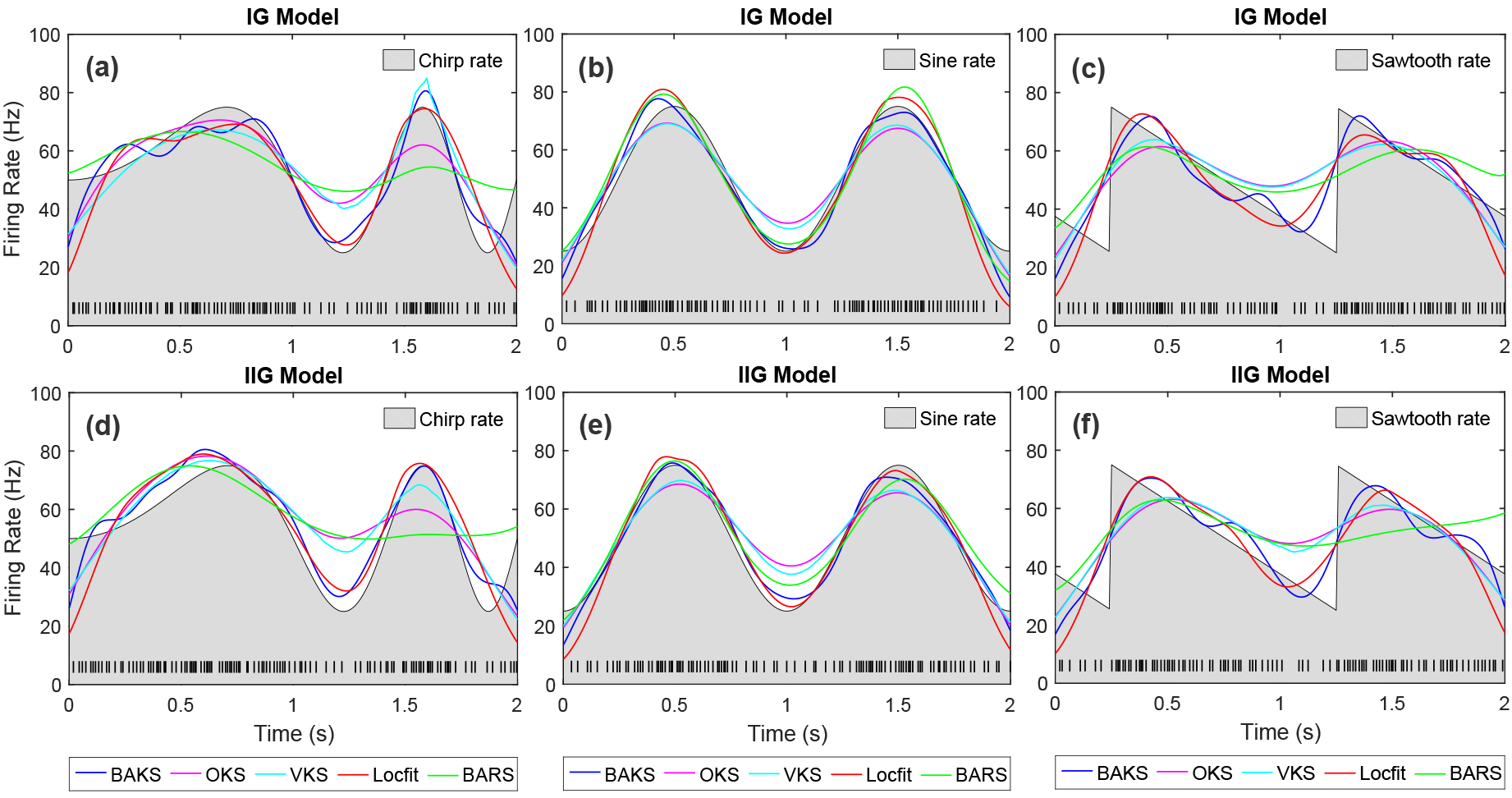
Comparison of firing rate estimates across all methods. (a)-(c) Firing rate estimates from IG model with chirp, sine, and sawtooth rate functions, respectively. (d)-(f) Firing rate estimates from IIG model with chirp, sine, and sawtooth rate functions, respectively. Black line plot with gray-shaded region indicates the underlying rate function. Black raster in the bottom of each plot represents the spike train generated from associated model and underlying rate function.

### 3.4. Comparison under different value of intensity and frequency

We studied the effect of different intensity and frequency of the underlying rate functions to the performance of the proposed method. In real neural data, the number of spikes (intensity) and the temporal fluctuation of spikes (frequency) may change slowly and rapidly. Therefore, we varied the intensity and frequency parameters as described in Table 1 (Testing 2). Using dataset testing 2, we performed 100 repetitions of single trial firing rate estimation from all methods. The MISE and its confidence interval comparison across all methods for the case of IG model is shown in Fig 6. From the total of 12 cases in IG model (3 rate functions and 4 variations of intensity and frequency), the BAKS method outperforms other competing methods in 9 cases (see Figs 6a, 6c and 6d). In the case of low frequency (Figs 6c), BARS shows better performance compared to the others. These results demonstrate the reliability of BAKS even when the intensity and frequency of underlying rate functions differ from that of during parameter tuning phase (training). Similar results are also observed for the cases of IIG model which can be seen in Appendix C (Fig C1).

### 3.5. Comparison under different value of shape parameter (γ)

The flexibility of the proposed method when the assumption of ISI shape deviates from that of used during the training phase is addressed in this section. Using dataset testing γ (γ = {1, 2, … 10}), we performed 100 repetitions of single trial firing rate estimation. From the total of 60 cases (2 models, 3 rate functions and 10 variations of γ value), 55 cases (91.67%) show the superior performance of the BAKS (blue triangle mark) over other methods as shown in Fig 7. In the 5 remaining cases, BAKS results in comparable performance to OKS and VKS methods which perform good under Poisson process (corresponds to γ = 1). These overall results demonstrate the flexibility of BAKS in estimating the firing rate from different ISI characteristic of spike train model.

### 3.6. Comparison under different number of trials

In offline analyses, the firing rate is typically estimated using spike trains from many similar trials aggregated into single compact spike trains. To study the effect of increasing number of trials, we assessed the performance of BAKS under different number of trials (*tr* = {5,10, 20, 30}) using dataset testing 4. We performed firing rate estimation using all methods, each for 100 times. The MISE comparison for this multi-trial cases is depicted in Fig 8. For all 6 scenarios, the increasing number of trials improves the performance of all methods as indicated by the decreasing MISE values. However, with the increasing number of trials, the rate of improvement declines as the MISE values reaching its convergence. Unlike in the single-trial case where the BAKS method outperforms all other methods, in some of multi-trial cases, the BAKS method cannot outperform all the others. For examples, in the cases of chirp and sine functions for both IG and IIG models, the BARS outperforms the BAKS method when *tr* ≥ 5 (Figs 8a, 8b and Figs 8d, 8e). In the case of sawtooth rate function, the BARS outperforms the BAKS on larger number of trials (*tr* = 30) as can be seen in Figs 8c and 8f. The OKS and VKS methods yield better estimates than that of BAKS only in case of sine rate function when *tr* ≥ 10 (Fig 8b and 8e). The performance of Locfit does not significantly improve when *tr* ≥ 5 and is relatively far behind of all other methods. As a summary, the BAKS method is the most superior in single-trial cases. In multi-trial cases with moderate number of spikes, the BAKS still shows good or comparable performance to other methods. Nevertheless, when it comes to multi-trial cases with large number of spikes, the BARS method performs the best among others.

**Figure 6.**
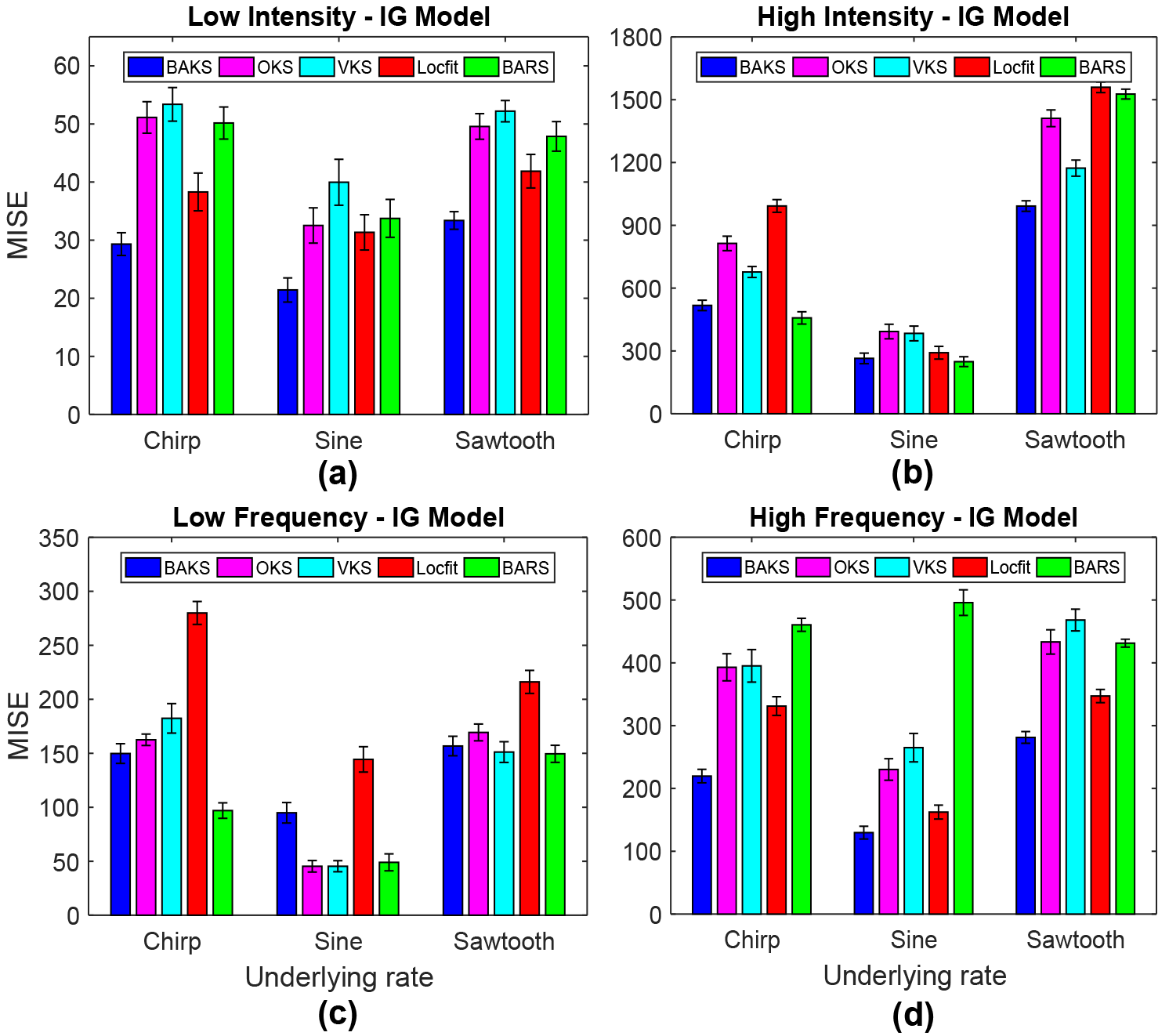
MISE comparison under different intensity and frequency for IG model. (a) MISE comparison for the case of low intensity. (b) MISE comparison for the case of high intensity. (c) MISE comparison for the case of low frequency. (d) MISE comparison for the case of high frequency. Vertical line crossing the peak of bar plot represents the 95% of MISE confidence interval.

**Figure 7.**
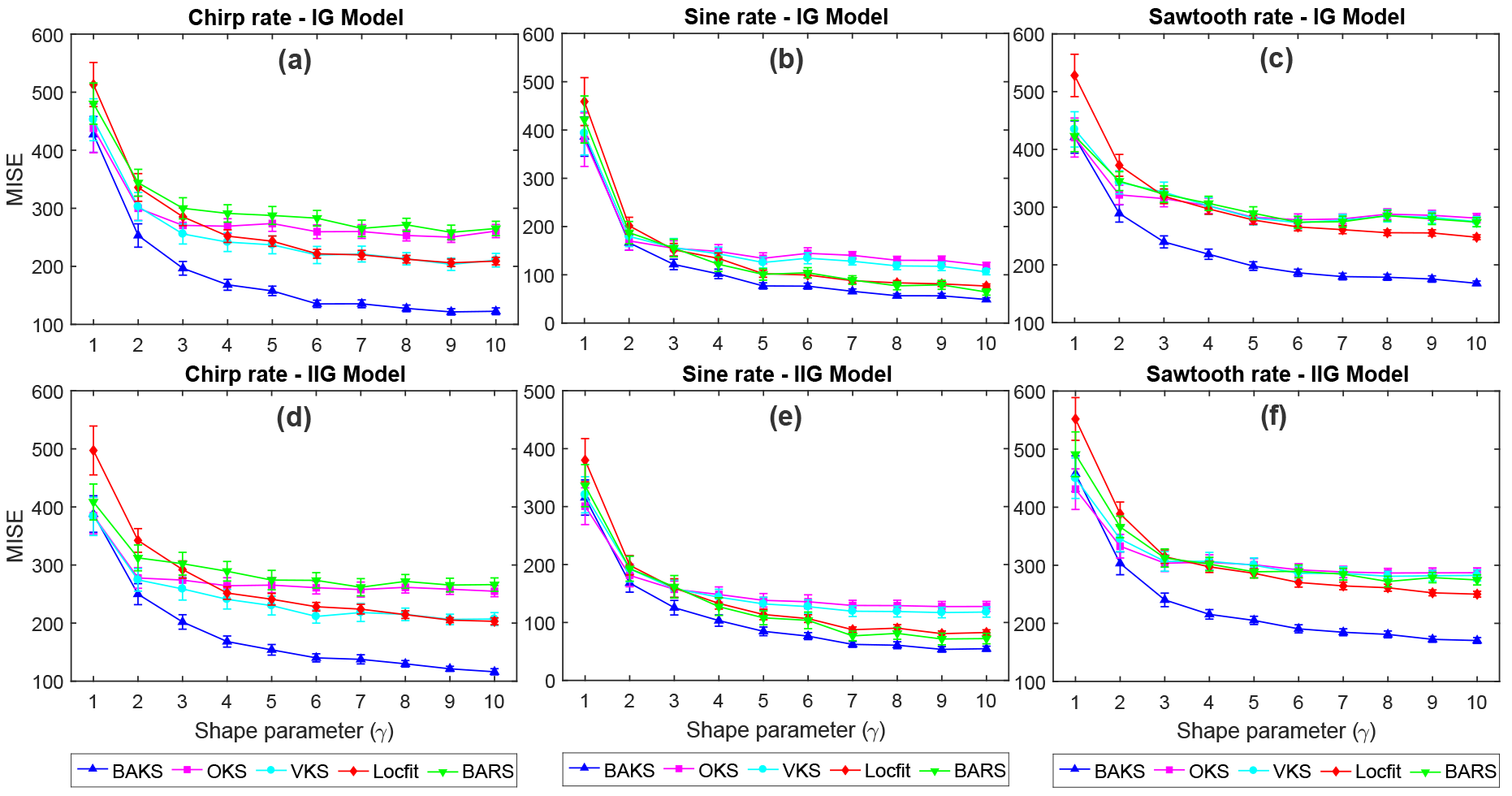
MISE comparison under various γ values. (a)-(c) MISE comparison for chirp, sine, and sawtooth rate functions, respectively, from IG model. (d)-(f) MISE comparison for chirp, sine, and sawtooth rate functions, respectively, from IIG model. Vertical bar represents the 95% of MISE confidence interval.

**Figure 8.**
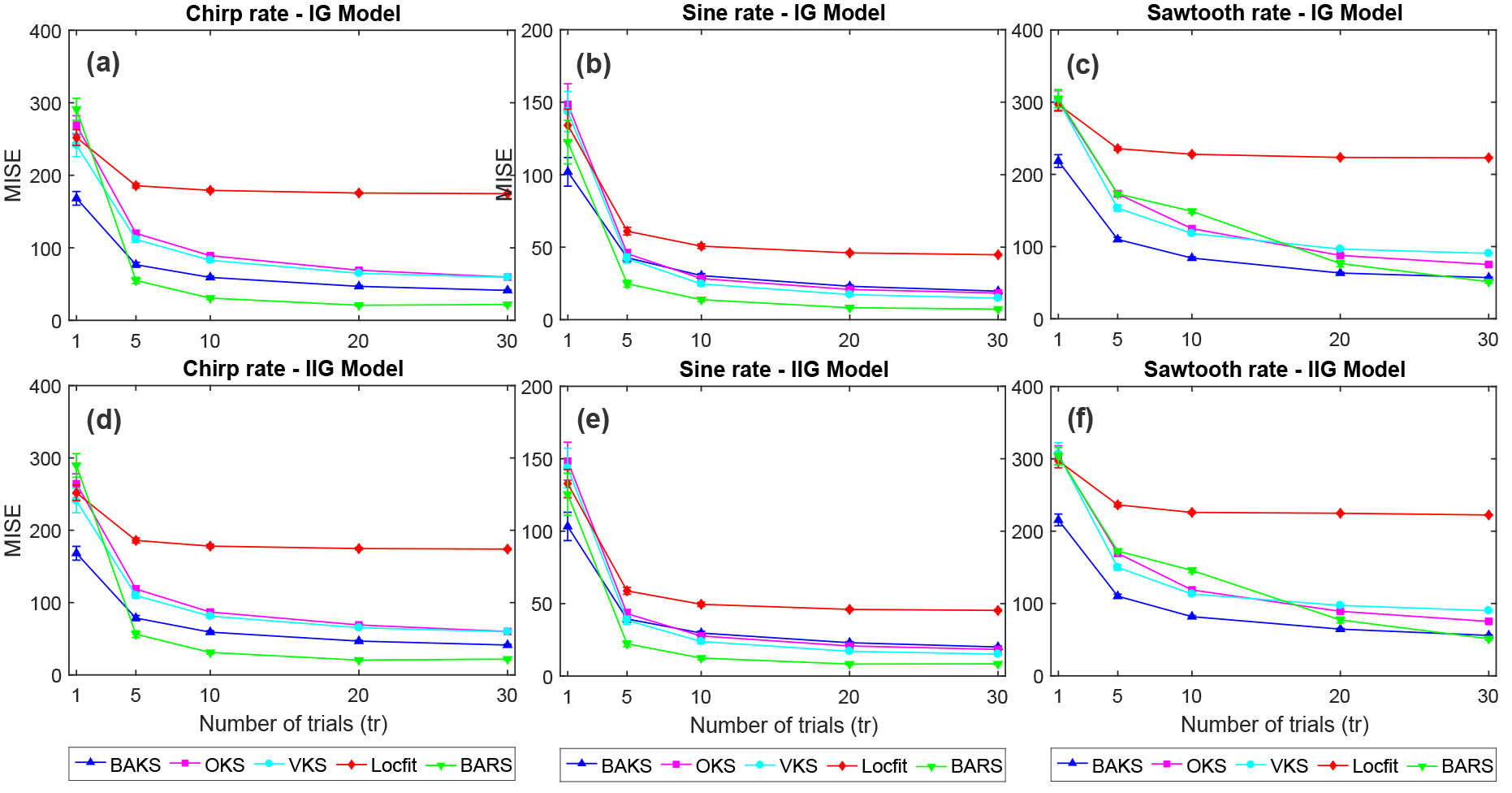
MISE comparison under different number of trials. (a)-(c) MISE comparison for chirp, sine and sawtooth rate function, respectively, for IG model. (d)-(f) MISE comparison for chirp, sine and sawtooth rate function, respectively, for IG model. Vertical bar (not clearly seen when *tr* ≥ 5) represents the 95% of MISE confidence interval.

### 3.7. Comparison under different underlying rate function

In practice, the true underlying rate function that generates the spiking data is unknown. There is infinite spaces of rate function that underlie the spiking generation. During the training, we used 3 underlying rate functions (chirp, sine and sawtooth) to find the optimal parameter for BAKS. Next, we studied the impact of different underlying rate function along with its intensity and frequency variations to the performance of BAKS. We performed 100 repetitions of firing rate estimation using dataset testing 5 which corresponds to single trial spike trains generated from Gaussian-damped sinusoidal rate function as in Eq (26). The performance comparison across all methods using this dataset for IG model is plotted in Fig 9. From a total of 6 cases (variation of intensity and frequency), BAKS outperforms other methods in 4 cases (all intensity cases and one medium frequency case). Similar results are also obtained for all the cases from IIG model which can be seen in Appendix C (Fig C2). This indicates the reliability of the BAKS even when the underlying processes depart from that of used during the training. The difference of underlying processes do not significantly affect the performance of BAKS.

**Figure 9.**
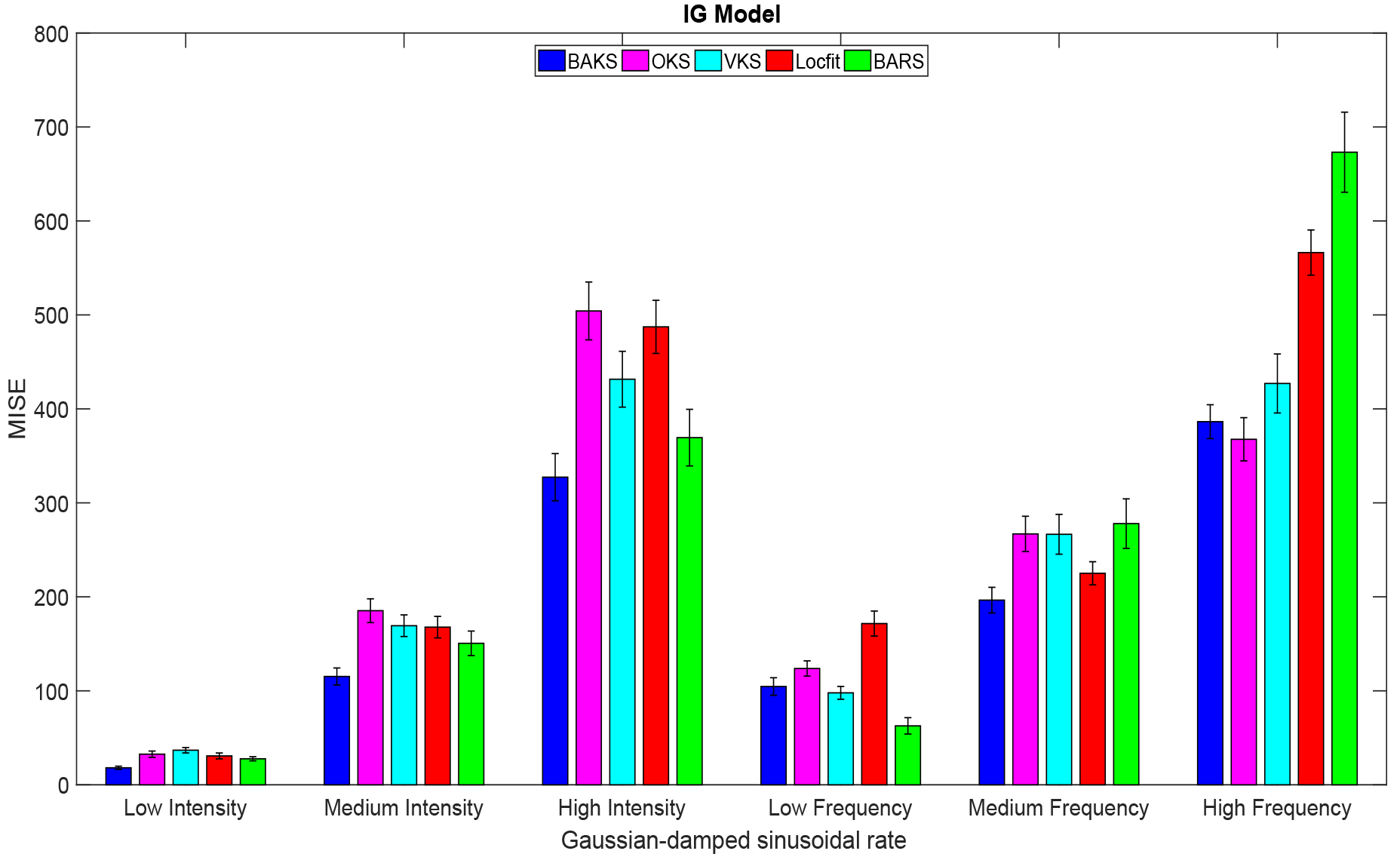
MISE comparison under different underlying rate function for IG model. Vertical line crossing the peak of the bar plot represents 95% of MISE confidence interval.

**Figure 10.**
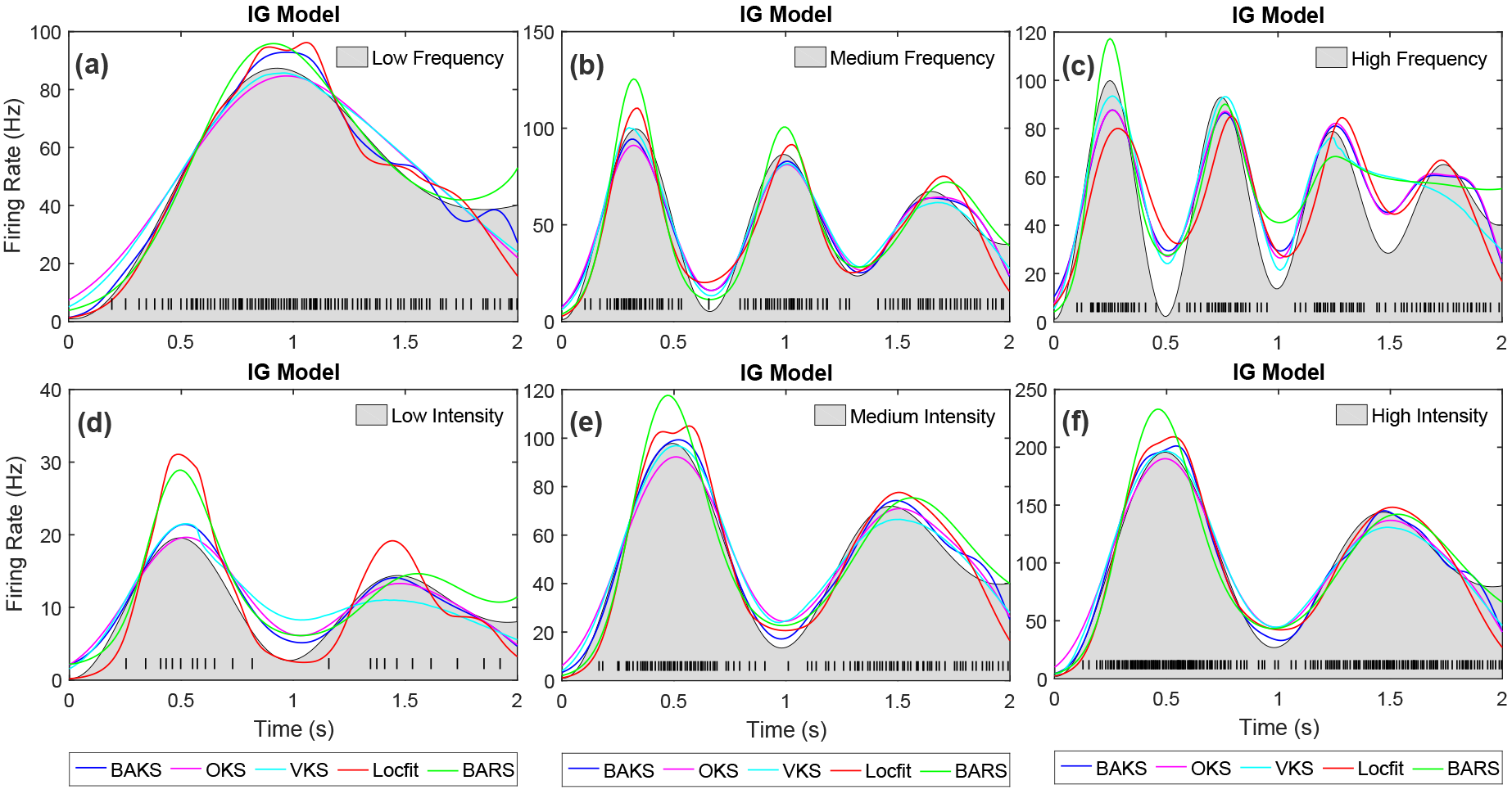
Firing rate estimate comparison for IG model with Gaussian-damped sinusoidal rate function. (a)-(c) Firing rate estimates for the cases of low, medium and high frequency, respectively. (d)-(f) Firing rate estimates for the cases of low, medium and high intensity, respectively. Black line plot with gray-shaded region indicates the underlying rate function. Black raster in the bottom of each plot represents the spike train generated associated with the underlying rate function.

The estimated firing rates from all methods across 6 scenarios for IG model with Gaussian sinusoidal rate function are shown in Fig 10. In all cases, BAKS method shows relatively consistent good visual fit. In the case of high frequency, BARS and VKS produce poor visual fit compared to the others (Fig 5c). In the case of low intensity, BARS and Locfit yield poor visual fit (Fig 10d) among others. In the remaining cases, the goodness-of-fit across all method does not differ significantly. Similar results are also observed for the cases from IIG model which is shown in Appendix C (Fig C3).

### 3.8. Comparison of computational complexity

The computational complexity reflects the execution time of a method. This is particular important when the above methods are applied on on-line firing rate estimation in BMI experiments in order to generate real-time feedback to the subject. This section describes and compares the computational complexity of each method.

Kernel smoothing technique has advantage of relatively simple and computationally fast. This is especially the case when the bandwidth is fixed throughout the observation interval such as in OKS method. OKS uses binned spike counts within certain bin interval (centered at estimation times) and kernel function with fixed bandwidth to estimate firing rate. The firing rate computation is performed by convolving the binned spike counts with the kernel function. OKS incorporates Fast Fourier Transform (FFT) for computing the convolution to further reduce the computation time. In OKS method, the bandwidth is selected by minimizing mean integrated squared error (MISE) function over the whole duration [12]. An extension to OKS method is VKS, which incorporates variable bandwidth. This variable bandwidth is computed by minimizing local MISE which requires large number of iterations with varying local interval. This iterative process makes VKS method significantly more complex than OKS. Unlike OKS and VKS, Locfit uses a polynomial to fit log-rate function by maximizing a local likelihood function. Locfit has relatively slow complexity because it uses fixed bandwidth selected in manual fashion; it does not employ automatic selection of bandwidth. Moreover, in this study, the bandwidth (in term of nearest neighbor) of Locfit was set to 0.4, meaning that the computation involves 40% of the total data within the whole duration.

Our proposed method, BAKS, even though incorporating automatic selection of optimal adaptive bandwidth, it still offers relatively low computational complexity. This advantage arises from the simple kernel smoothing technique with proper choice of prior distribution and kernel function which leads to closed-form expression of posterior bandwidth. This in turn simplifies the computation process of determining the adaptive bandwidth. This type of convenient closed-form expression cannot be obtained in the case of BARS method; thus, a numerical approximation technique is required. BARS uses iterative procedure involving computationally expensive Markov chain Monte Carlo (MCMC) technique and Bayes information criterion (BIC) to find the optimal smoothing parameters. This process takes relatively long computation to yield “converged” results.

The computational complexity among these methods can be indicated by the time required for completing the firing rate estimation in computer simulation (i.e. computational runtime). Since this runtime comparison is impacted by the code implementation of each method, this should be viewed as estimation of the real computational complexity of each method. The code for BAKS can be downloaded from https://github.com/nurahmadi/BAKS. The OKS and VKS codes can be downloaded from http://www.neuralengine.org/res/kernel.html. The Locfit code is available through http://chronux.org/; while the BARS code is available through http://lib.stat.cmu.edu/~kass/bars/bars.html. All the program codes were written and run in Matlab R2016b software (The Mathworks Inc., Natick, MA) installed on Windows 7 64-bit PC with 8 Intel cores i7-4790@3.6 GHz and RAM 16 GB. This comparison uses 100 repetitions of single trial synthetic spike train data (2s duration) generated by IG model with chirp rate function.

As shown in Table 2, Locfit and OKS are the two fastest methods; both methods complete the computation within the order of few milliseconds. VKS, on the other hand, requires around 3 order of magnitude longer time than both Locfit and OKS; whereas BARS method requires longest time (in the order of seconds). In the BARS parameter setting, we set burn-in iterations to 500 and sample iterations to 5000. The BARS’ runtime can be reduced by setting the burn-in and sample iterations to smaller values. However, this may result in decreasing accuracy as the trade-off. Therefore, these parameter should be carefully set to find good trade-off between accuracy and runtime. Wallstrom suggested that the default values for burn-in and sample iterations are 500 and 2000, respectively [40].

**Table 2.** Average runtime comparison of BAKS with other methods (in second)

| Single trial case | BAKS | OKS | VKS | Locfit | BARS |
| --- | --- | --- | --- | --- | --- |
| Low intensity | 0.0016 | 0.0017 | 0.2340 | 0.0012 | 4.8438 |
| Medium intensity | 0.0062 | 0.0018 | 0.3306 | 0.0012 | 4.6563 |
| High intensity | 0.0126 | 0.0017 | 0.3306 | 0.0014 | 4.5527 |

The runtime performance of BAKS is significantly better than both VKS and BARS, and is slightly below that of Locfit and OKS. Table 2 shows that BAKS’ runtime is influenced by the intensity of the underlying rate function (i.e. number of spikes), whereas other methods’ runtime are relatively consistent. This is because other methods incorporate binning procedure for the spiking data prior to their core computation. This makes the number of input data fed to the core computation always uniform regardless of the number spikes within the observation interval. This is not the case for BAKS method. Our current BAKS code is a straightforward implementation of the formula described in Methods section. In this study, we have not considered the efficient implementation of BAKS method. It is important to note, as neurons have a property of refractory period, the number of spikes within observation interval is limited. This guarantees that under single trial, even with current implementation code, the runtime performance of BAKS will only decrease up to certain bound.

### 3.9. Application to real neural data

In this section, we apply our proposed method for estimating the neuronal firing rate from real data obtained from two public neural databases, which are database for reaching experiments and models (DREAM) and neural signal archive (NSA). The DREAM and NSA databases can be accessed from http://crcns.org/ and http://www.neuralsignal.org/, respectively. In the DREAM database, we used Flint_2012 dataset that were recorded from primary motor cortex area (M1) of monkey’s brain when the subject was performing center-out reaching task. Single unit spikes were obtained by using thresholding and offline sorting technique. More detailed information on the recording tools and experimental setup can be found in [41]. In the NSA database, we used nsa2004.1 dataset recorded from visual cortex (MT/V5) area when random dot stimuli was being presented to a monkey [42]. The detailed electrophysiological recording is given in [43].

Unlike the synthetic data in which the true underlying rate function is known, in the case of real neural data, we do not have access to the underlying rate (i.e. ground truth). Therefore, the ‘true’ underlying rate in real neural data was estimated by averaging and smoothing the spike counts across many similar trials. These similar trials were selected such that each trial contains greater or equal to 50 spikes/s within observation interval (1s for the Flint_2012 dataset, 2s for the nsa2004.1 dataset). This limited number of spikes was taken on the assumption that neurons likely fire more spikes when performing tasks or receiving stimuli. In this work, We considered only neurons that satisfy this criterion in more than 30 trials in order to obtain sufficiently small error as we observed in multi-trial synthetic data (Fig 8). To this end, we obtained 47 (4) subdatasets with total trial of 1791 (134) for the Flint_2012 (nsa2004.1) dataset. In the Flint_2012 dataset, we aligned the spiking responses over same-direction reaching tasks to the time when the monkey started the actual hand movement (indicated by cursor movement). To make the observation interval the same from inherently different trial duration for each trial, we used on average 200ms before and 800ms after the movement (total duration 1s). In the nsa2004.1 dataset, the spiking responses were aligned to the time when random moving dot stimuli was firstly presented to the monkey. In this type of experiment, the trial duration was fixed to 2 s.

To estimate the ‘true’ underlying rate, we first superimposed the spike trains from all trials of one neuron. We partitioned the observed duration into 10ms bin interval and computed the spike count within the bin interval. The ‘true’ underlying rate was then smoothed by using BARS method since the BARS demonstrates the most superior performance on the synthetic data when the number of trial is large (*tr* = 30, see Fig 8). However, in this multi-trial case where there exists spike train variability across trials, we increased the sample iteration to 30,000 and burn-in samples to 5,000 to ensure the convergence of BARS results. One of the advantages of the BARS is that it provides the output estimate along with its confidence interval. In this work, we used 95% confidence interval. When computing MISE, we took into account this uncertainty. Before squaring and integrating across observation interval, we normalized the error between ‘true’ underlying rate obtained from many trials (act as a reference) and singletrial estimated firing rate from method of interest by dividing it with upper or lower confidence interval. The upper (lower) interval was used when the estimated firing rate is larger (smaller) than the reference. By doing so, we impose more (less) weight when the confidence interval is smaller (larger) to adjust the uncertainty brought by the BARS estimation. We call this normalized MISE as a weighted MISE (WMISE) and formulate it as follows,

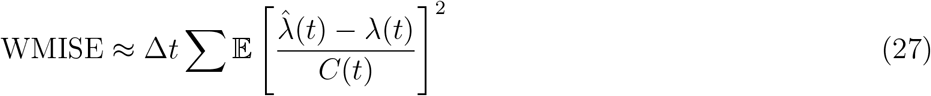

where *C*(*t*) is set to the upper confidence interval when 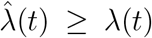 and the lower confidence interval when 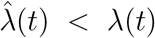. These upper and lower confidence intervals calculated by the BARS are not uniform.

We examined the performance comparison across methods under WMISE function. Based on WMISE function, we derived three different metrics for performing the comparison. First, we investigated the average WMISE performance across total number of trials. Based on 1791 single-trial firing rate estimation from 47 subdatasets in the Flint_2012 dataset, the BAKS method produces the smallest WMISE average and confidence interval (10.34 ± 0.53) as shown in Fig 11a. The BAKS method also produces the best performance (19.77 ± 2.45) in the case of nsa2004.1 dataset with total 134 trials from 4 subdatasets (Fig 11d). Second, we measured the number of times (in percentage form) one method outperformed all the other methods. In both Flint_2012 and nsa2004.1 datasets, the BAKS more frequently (41.99% and 42.54% respectively) outperforms the other methods as shown in Figs 11b and 11e. In these cases, the OKS method comes as the second with 30.65% and 17.16%. Third, we assessed the average improvement (WMISE decrease in %) per trial of the BAKS method over other methods. Figs 11c and 11f describes the BAKS performance compared to other methods for the Flint_2012 and nsa2004.1 datasets. A positive (negative) bar value means that the BAKS method outperforms (is outperformed by) the others. As described in Fig 11c, the single-trial average improvement of the OKS method is slightly better than the BAKS (-2.58 ± 1.90%). This seems to contradict the results when using the first and second metrics, in which the BAKS method outperforms the OKS. By plotting the percentage of average improvement of the BAKS against the OKS across 1791 trials, it turns out that although the OKS fewer times outperforms the BAKS (than that of the opposite), but in few trials the magnitude of improvement made by the OKS is larger than the BAKS. Contrary to the Flint_2012 dataset, the BAKS method is superior to the OKS (5.37 ± 3.75%) in the nsa2004.1 dataset (Fig 11f). This is consistent with the results measured by the two other metrics. On average, the BAKS method consistently performs good compared to other methods in both datasets. Some examples of single-trial firing rate estimation from all methods for the Flint_2012 and nsa2004.1 datasets are shown in Fig 12. Figs 12a and 12b show the firing rate estimates from ‘N184’ and ‘E164’ cases in the Flint_2012 dataset. Figs 12c and 12d show the firing rate estimates from ‘j032_25.6’ and ‘j032_51.2’ cases in the nsa2004.1 dataset. The firing rate estimates from BAKS are visually comparable to other methods. However, BARS show poor visual fit in towards the boundaries (beginning or end of the underlying rate function).

**Figure 11.**
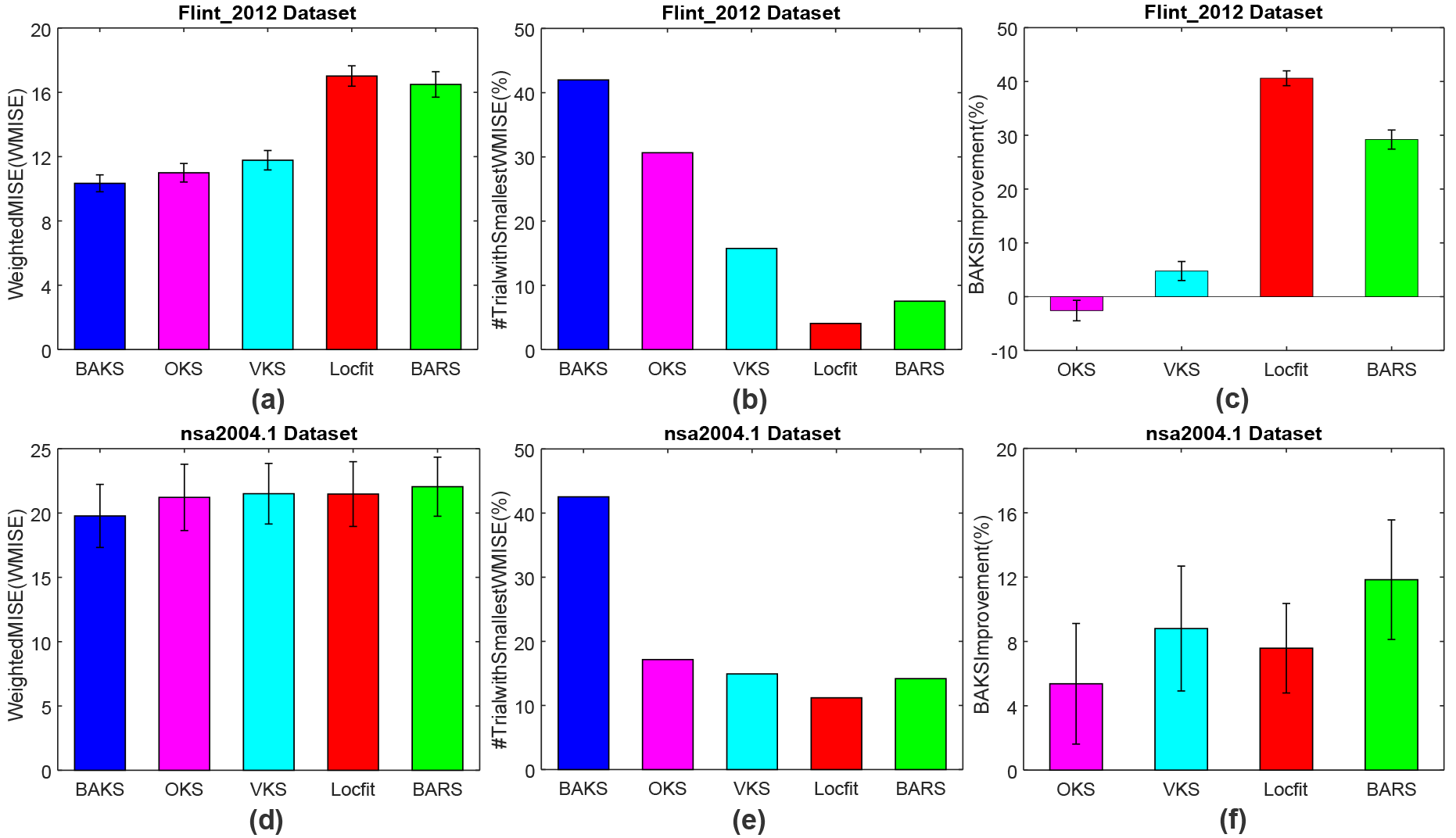
WMISE comparison across all methods using real neural data. (a) Average WMISE comparison across all trials in Flint_2012 dataset. (b) Number of times (in %) BAKS outperforms other methods in Flint_2012 datasets. (c) Single trial performance improvement made by BAKS over other methods in Flint_2012 datasets. (d)-(f) Similar to that of (a)-(c) but with nsa2004.1 dataset. Vertical crossing the peak of the bar plot represents the 95% of WMISE confidence interval.

**Figure 12.**
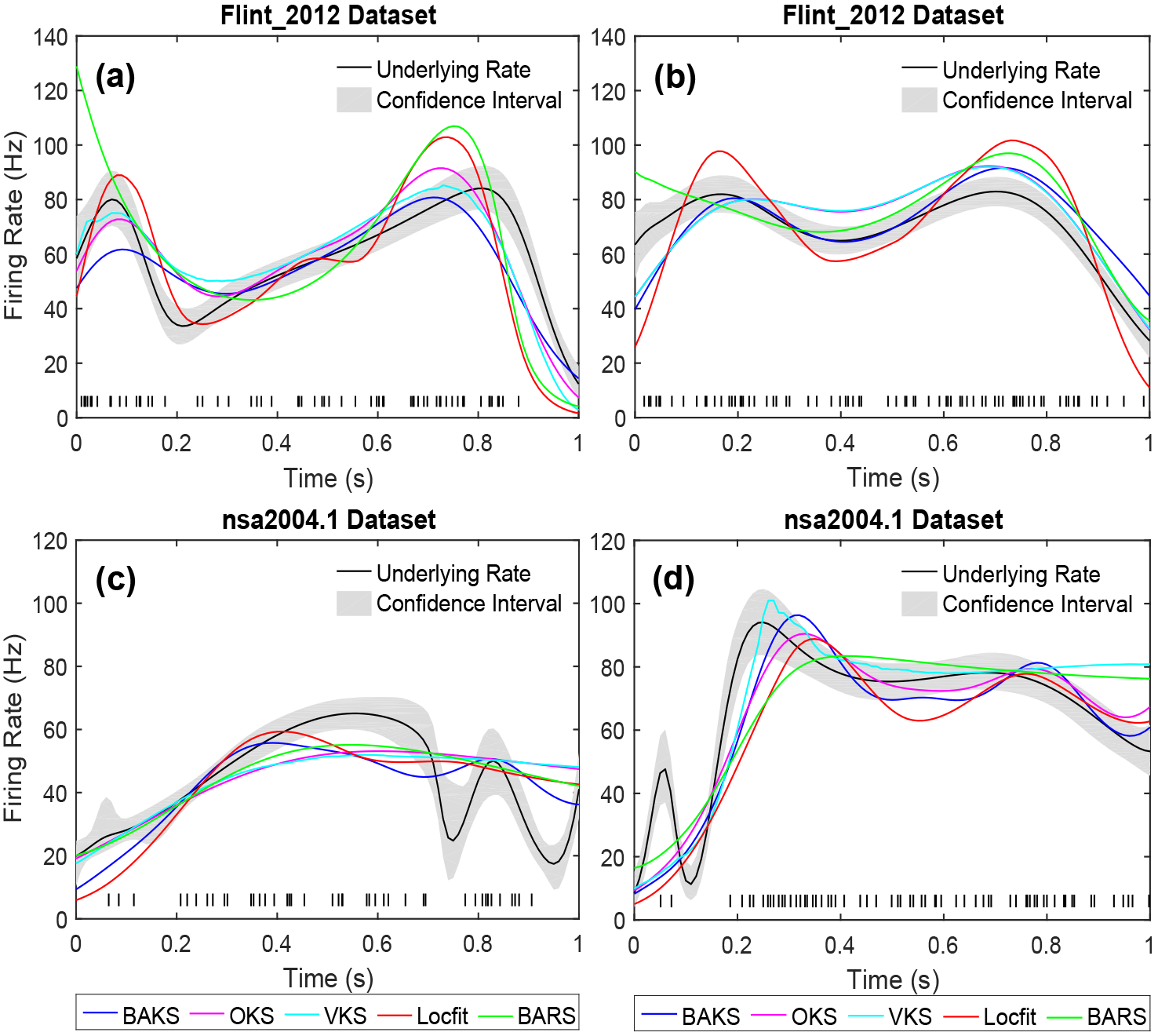
Firing rate estimates comparison for real neural data. (a) Firing rate estimates from ‘N184’ of Flint_2012 dataset (b) Firing rate estimates from ‘E164’ of Flint_2012 dataset (c) Firing rate estimates from ‘j032_25.6’ of nsa2004.1 dataset. (d) Firing rate estimates from ‘j032_51.2’ of nsa2004.1 dataset. Black line plot and gray-shaded area indicate the ‘true’ underlying rate (estimated from ≥ 30 trials) and its 95% confidence interval, respectively. Black raster in the bottom of each plot represents the single spike train taken from the associated underlying rate.

## 4. Discussion

In this study, we propose a new method for estimating single-trial neuronal firing rate. Our method employs a kernel smoothing technique with adaptive bandwidth. This differs from other kernel-based firing rate estimation methods in that its bandwidth parameter is adaptively determined by a Bayesian approach. The proposed method, BAKS, has been developed with the motivation to estimate firing rate from single spike train generated from underlying rate function that dynamically changes over observed duration (non-stationary).

We select the optimal parameter of BAKS using a synthetic spike train stochastically sampled from 3 rate functions (as representation of non-stationary underlying processes). These rate functions are chirp, sine, and sawtooth expressed in Eqs (23), (24) and (25), respectively. Using this optimal parameter, we evaluate the performance of BAKS using 5 synthetic datasets. These datasets represent various setting and combination of underlying rate functions along with their intensity and frequency variations, ISI shape (parameter γ), and number of trials. The performance comparison is measured under MISE function. By extensive simulations, we demonstrate good performance of the proposed method, BAKS, compared to two other kernel-based methods (OKS and VKS) and two generalized nonparametric regression methods (Locfit and BARS). On average, BAKS outperforms the other methods in single-trial estimation (smallest MISE) across various settings. The adaptive bandwidth featured in the BAKS can adjust the different spike densities within the observation interval. The results suggest that our proposed BAKS method is suitable to be used for single-trial analysis of neural data. The flexibility of the BAKS has also been tested by using spike train generated from the same 6 scenarios but with different shape parameter values (γ = {1, 2, … 10}). Consistent results are obtained despite using these different characteristics of spike train. The BAKS method does not assume specific distribution on the spike train, rather it uses appropriately chosen prior distribution on the bandwidth parameter. The prior distribution of the bandwidth is derived from Gamma prior distribution on the precision parameter (inverse of square bandwidth). The precision parameter describes how concentrated observed are around the means of Gaussian kernel which are set to the spike times. Since these spike times (i.e. sum of ISI) are conveniently modeled with Gamma distribution [15,26–29], the precision parameter is also assumed to be Gamma distribution. This choice has been shown to yield good performance. On the other hand, all other competing methods (OKS, VKS, Locfit, and BARS) use a Poisson assumption [12–14], which is less likely for the case of single-trial spike train; neurons have certain properties (e.g. refractory and bursting) that cannot be described by the Poisson model [15,16]. Numerous works have shown the inadequacy of the Poisson model and proposed other more biophysically plausible models (e.g. IG and IIG) [6,15,26–29,34]. The deviation from Poisson assumption under single-trial cases may lead OKS, VKS, Locfit and BARS to poor performance [12,17].

We also compare the performance of the BAKS method under multi-trial cases to study the implication of increasing number of spikes within the same duration. Some topics of interest in neuroscience use multi-trial spike train to obtain the firing rate. The results show that all methods produce similar trend; the increasing number of trials up to a certain value (threshold) improves the performance (smaller MISE). However, as the number of trial increases, the rate of improvement of each method and the threshold value differ from each other. The BARS is the one that improves performance significantly with increasing number of trials. It has been suggested that under many trials, superimposed spike train across trials can be approximated by inhomogeneous Poisson model [12,16]. With a sufficient number of spike counts within the bin intervals (approximating Poisson count) together with intensive computation in determining optimal knot configurations, the performance of BARS improves significantly. In the case of chirp and sine rate functions, the BARS starts to outperform the BAKS method within few number of trials (*tr* = 5). However, in the case of sawtooth rate function, the BARS needs larger number of trials to outperform the BAKS (*tr* = 30). Nevertheless, in comparison to OKS, VKS, and Locfit methods, the BAKS method still yields relatively good performance. The overall results suggest that the BAKS method is good at estimating firing rate from a low to moderate number of spikes (represented by single or few trials) from non-stationary underlying rate functions. The BAKS performance is especially good in the case of discontinuous rate functions (e.g. sawtooth).

After validation using synthetic data, the proposed BAKS method is also tested using real neural data recorded from motor and visual cortex of non-human primate (NHP). The motor neural data (Flint_2012) is associated with center-out reaching tasks, whereas the visual neural data (nsa2004.1) is associated with moving random-dot visual stimuli. Measuring the performance in real neural data is a challenging due to unknown underlying rate. Hence, the underlying rate is estimated by using multi-trial cases on the assumption that neurons respond similarly upon given similar tasks/stimuli. This procedure is similar to the one described in [26]. However, in practice, neuronal response may considerably differ across similar trials. To minimize large variation in the spike trains, subsets from two datasets (Flint_2012 and nsa2004.1) are selected with constraints explained in previous section. The BARS method is chosen to estimate the underlying rate as it provides the smallest MISE in multi-trial synthetic data. To account for the estimation uncertainty produced by the BARS, weighted MISE (WMISE) is used for comparison. Among three metrics and two real datasets (6 cases) that we use, there is only 1 case that one other method (OKS) outperforms the BAKS (see Fig 11c). The performance improvement of the OKS over the BAKS in this only case is relatively small (2.58%). The overall results show that, on average, the BAKS method yields good performance compared to all other competing methods. This is in good agreement with the results obtained from single-trial synthetic data, which further demonstrates the effectiveness of the proposed method in estimating single-trial neuronal firing rate.

The BAKS method offers ease and simplicity as standard kernel-based method does, yet effective in grasping sudden and slow changes of firing rate in different region within the observation interval. Unlike the BARS method which is computationally demanding, the BAKS method is relatively fast owing to analytical expression of bandwidth posterior density. This analytical expression leads to the adaptive bandwidth determined in an exact way (not numerical approximation), which reduces the computational complexity. With good performance and relatively low complexity, BAKS is suitable to be used for research that require single-trial firing rate estimation. For examples, understanding the encoding mechanism of neurons in cognitive-related tasks and decoding task parameter in brain-machine interface (BMI) applications. As a summary, the comparison of BAKS with other methods is given in Table 3.

**Table 3.** Comparison summary of BAKS with other methods.

|  | BAKS | OKS | VKS | Locfit | BARS |
| --- | --- | --- | --- | --- | --- |
| Adaptability to underlying dynamics | ✓ | — | ✓ | — | ✓ |
| Bayesian/probabilistic approach | ✓ | — | — | — | ✓ |
| Automatic selection of smoothing parameter | ✓ | ✓ | ✓ | — | ✓ |
| Single trial (low to moderate number of spikes) | ◇◇◇ | ◇◇ | ◇◇ | ◇◇ | ◇◇ |
| Multi trials (large number of spikes) | ◇◇ | ◇◇ | ◇◇ | ◇ | ◇◇◇ |
| Computational complexity (runtime) | ◇◇◇ | ◇◇◇ | ◇◇ | ◇◇◇ | ◇ |

More diamonds mark (◇) indicates better (desirable) performance.

## 5. Conclusion

We have presented a simple yet accurate method for estimating single-trial neuronal firing rate based on kernel smoothing technique with adaptive bandwidth. The key idea of this method is to consider the bandwidth parameter as random variable under a Bayesian framework. By using Bayes’ theorem with proper choice of kernel and prior distribution functions, the bandwidth can be adaptively determined in an exact and quick way. Extensive evaluations on both synthetic and real neural data show that the proposed method yields good performance compared to other competing methods. This suggests that the proposed method has the potential to improve single-trial analysis in neuroscience studies and decoding performance of spike-based brain-machine interfaces (BMIs).

## Appendix A. Posterior distribution of bandwidth

In this appendix, we derive a closed-form expression of the posterior density of bandwidth as given in Eq (10). According to Bayes’ theorem, the posterior density is formulated as:

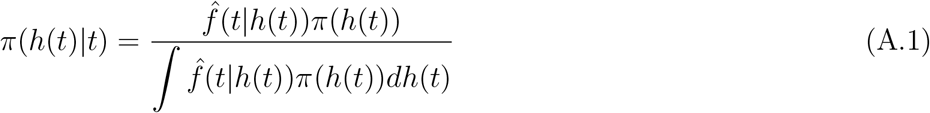

By using likelihood function as in Eq (8) and prior distribution of bandwidth as in Eq (7), we can obtain:

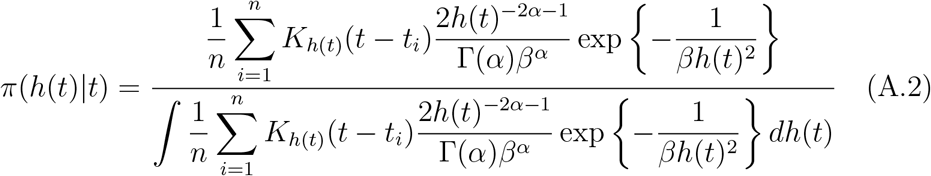

By substituting Gaussian kernel into the likelihood function and removing the same constants in both numerator and denominator, Eq (A.2) then becomes:

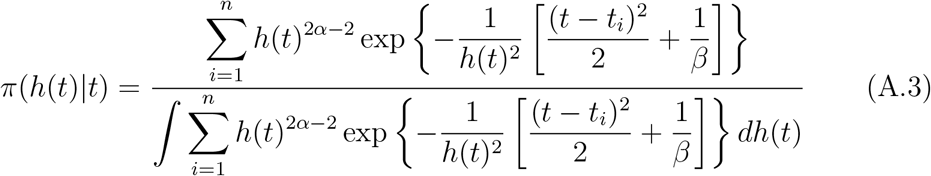

Let now consider the denominator of Eqs (A.1) and (A.3), which we can rewrite as:

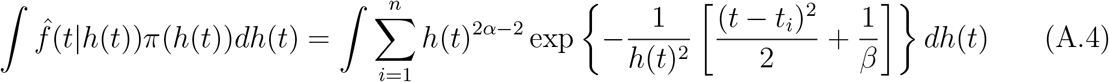

To simplify the calculation, let us define variables as follows:

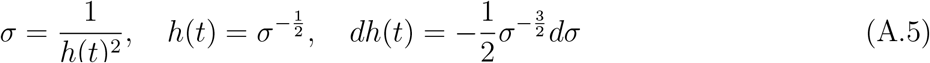

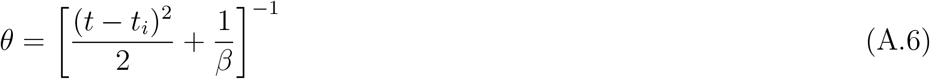

By substituting Eqs (A.5) and (A.6) into Eq (A.4), the integral function can be represented as:

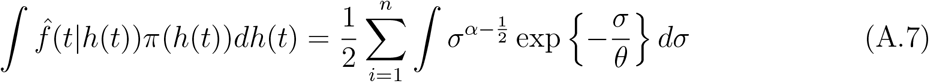

Eq (A.7) can be simplified so that the integral part forms Gamma probability density as follows:

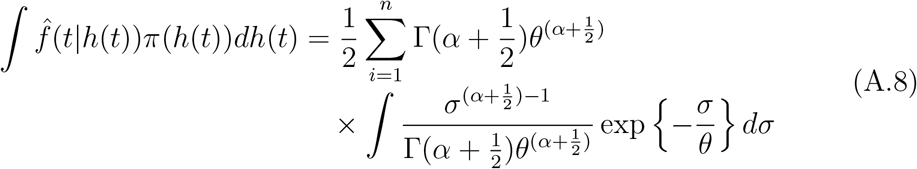

Since the integration of Gamma probability density function is equal to 1,

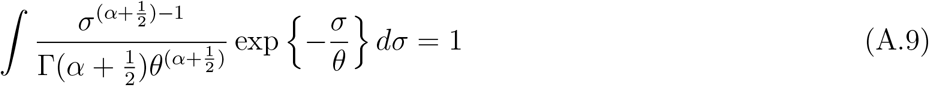

Eq A.8 can then be analytically expressed as:

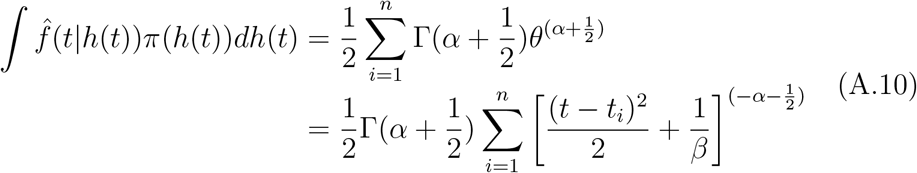

Finally, by substituting Eq (A.10) back to the original equation of posterior density of bandwidth in Eq (A.1), we can obtain the closed-form solution as in Eq (10):

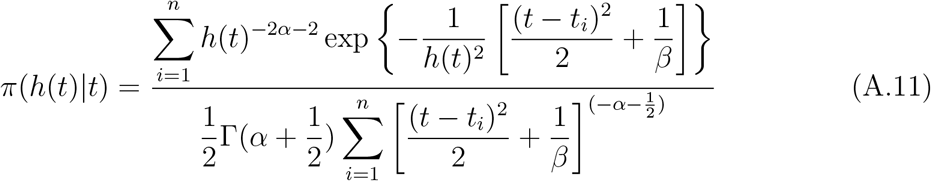

## Appendix B. Adaptive bandwidth estimate

Under squared error loss function, the adaptive bandwidth can be estimated by using the posterior mean as given by:

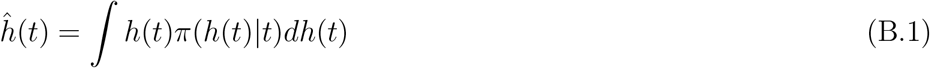

By substituting Eq (A.11) into Eq (B.1), we can obtain:

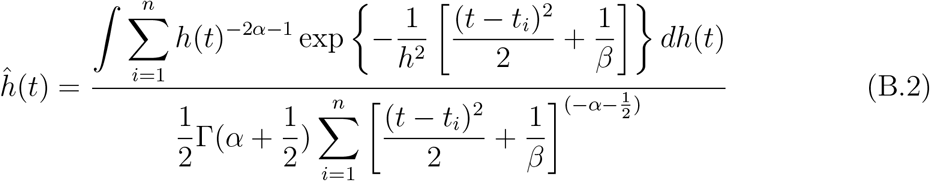

Similar to the derivation procedure for the posterior distribution of bandwidth (Appendix A), by the change-of-variables rule using Eqs (A.5) and (A.6), Eq (B.2) can be written as:

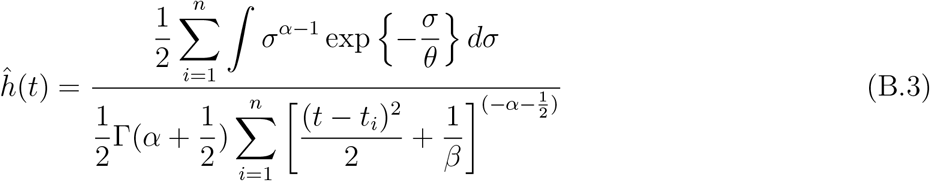

By modifying the integral part of numerator to be an integration of Gamma probability density function (which is equal to 1) as in Eq (A.8), we can obtain the final closed-form solution as in Eq (12):

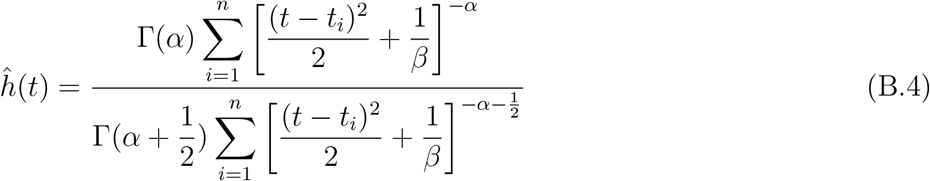

## Appendix C. Supporting information

**Figure C1.**
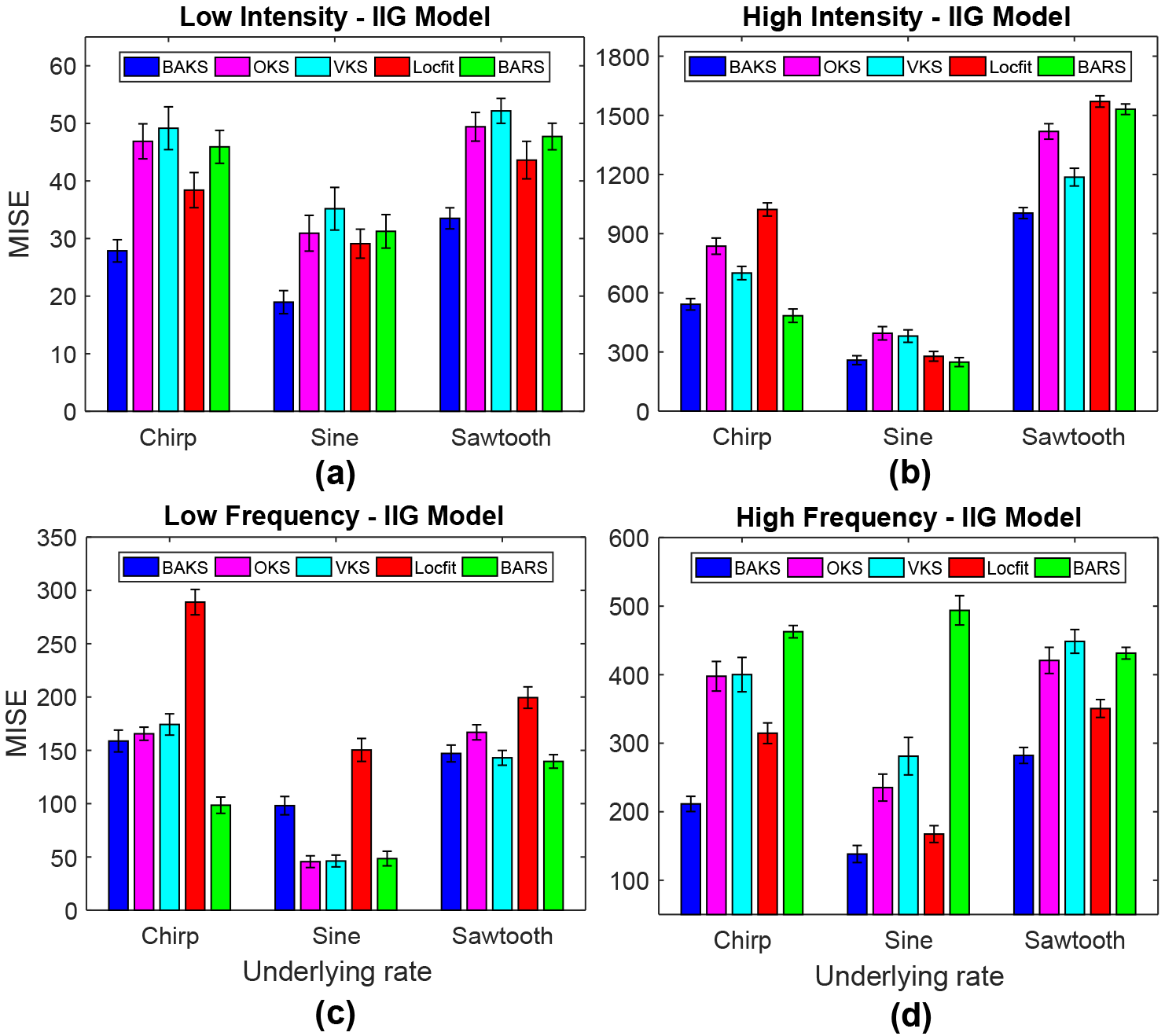
MISE comparison under different intensity and frequency for IIG model. (a) MISE comparison for the case of low intensity. (b) MISE comparison for the case of high intensity. (c) MISE comparison for the case of low frequency. (d) MISE comparison for the case of high frequency. Vertical line crossing the peak of bar plot represents the 95% of MISE confidence interval.

**Figure C2.**
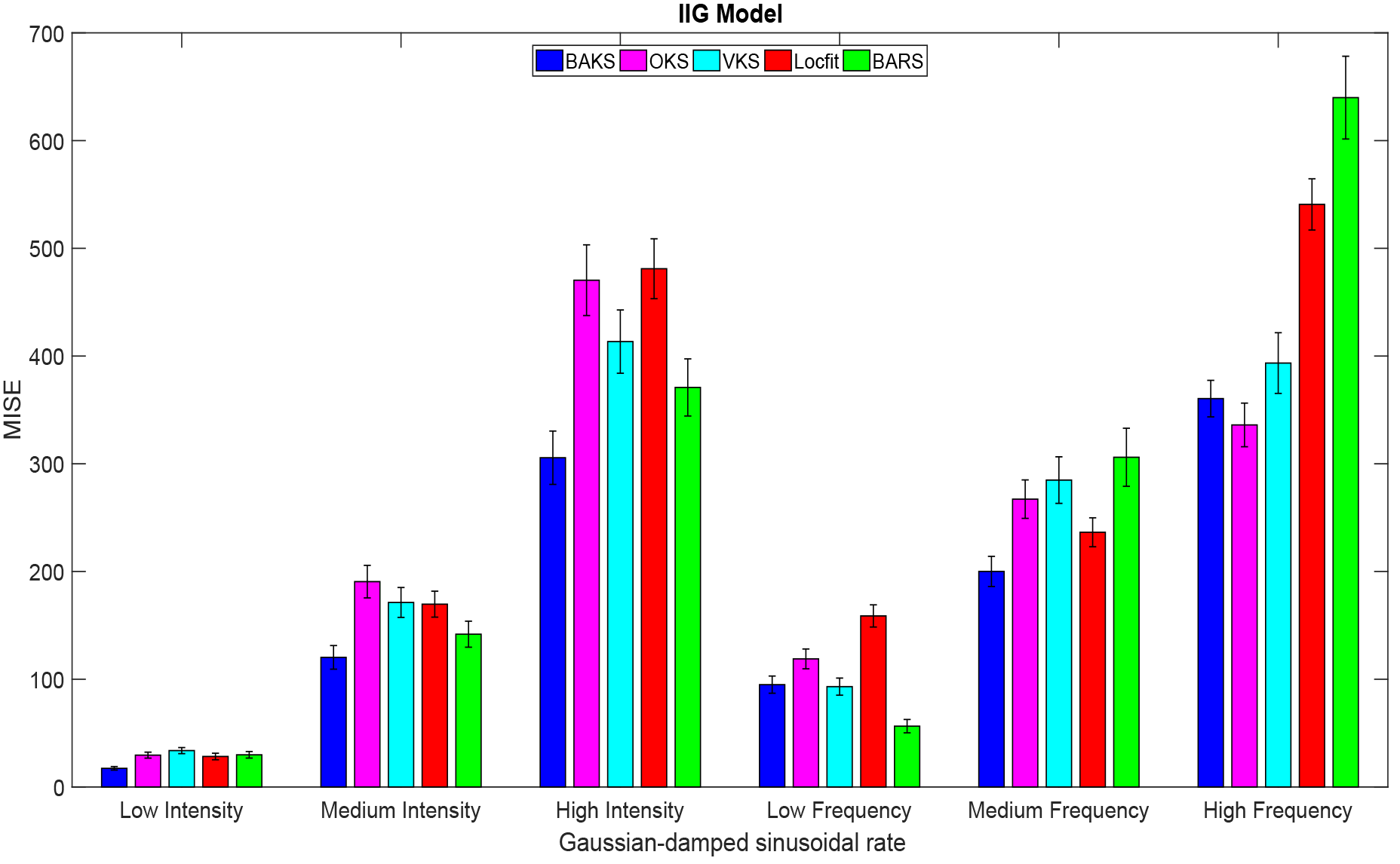
MISE comparison under different intensity and frequency for IIG model. Vertical line crossing the peak of the bar plot represents 95% of MISE confidence interval.

**Figure C3.**
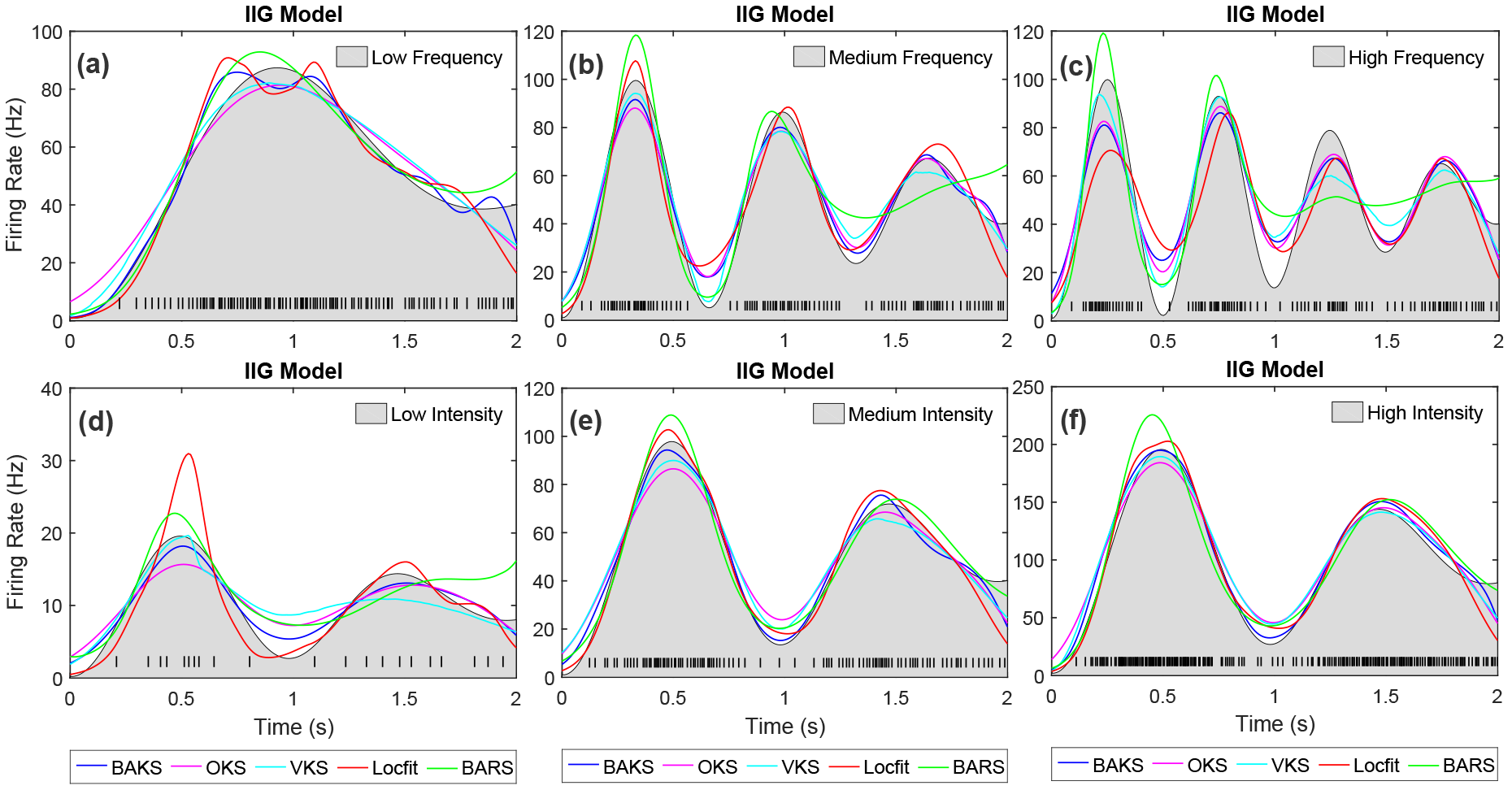
Firing rate estimate comparison for IIG model with Gaussian-damped sinusoidal rate function. (a)-(c) Firing rate estimates for the cases of low, medium and high frequency, respectively. (d)-(f) Firing rate estimates for the cases of low, medium and high intensity, respectively. Black line plot with gray-shaded region indicates the underlying rate function. Black raster in the bottom of each plot represents the spike train generated associated with the underlying rate function.

## Acknowledgments

We thank R.D. Flint, E.W. Lindberg, L.R. Jordan, L.E. Miller, and M.W. Slutzky for making their data available via the CRCNS database (https://crcns.org/). We also thank K.H. Britten, M.N. Shadlen, W.T. Newsome, and J.A. Movshon for making their data available via the NSA website (http://www.neuralsignal.org/). Nur Ahmadi acknowledges the graduate scholarship granted by Indonesia Endowment Fund for Education (LPDP), Republic of Indonesia.

